# Metabolic cascades drive spatial prebiotic fermentation in a defined gut consortium

**DOI:** 10.64898/2026.10.30.763039

**Authors:** Dorothée L. Berthold, Nicola R. Realini, Andreas Sichert, Annika Hausmann, Uwe Sauer

## Abstract

Dietary fiber deficiency shifts the gut microbiota to degrade host mucus, compromising intestinal homeostasis. Mechanistically linking *in vitro* metabolic dynamics of fiber degradation to *in vivo* outcomes remains challenging due to a lack of representative model systems. Introducing spatial mucin structures to a defined Oligo-Mouse-Microbiota (OMM12) *in vitro* consortium allowed us to recapitulate the spatio-temporal dynamics of xylooligosaccharide (XOS) processing in gnotobiotic mice. XOS supplementation rapidly established a stable community state *in vitro* and *in vivo* across compositional, metabolic and transcriptional levels. XOS supplementation established distinct functional guilds: *Bifidobacterium animalis* and *Blautia coccoides* partitioning xylose and arabinose catabolism, whilst profiting from XOS degradation by *Bacteroides caecimuris*. Along the colonic axis, a spatial XOS gradient exposes *Bl. coccoides*’s substrate preference for xylose and drives longitudinal functional adaptations. Altogether, our predictive platform proves that regional microenvironments govern prebiotic efficacy, highlighting spatial cascades as essential parameters for targeted microbiome interventions.

## Introduction

The mammalian gut microbiome assembles through an orchestrated hierarchy where host habitat filtering, dietary inputs, priority effects and bacterial competition govern niche availability across ecological guilds^1,2^. However, defining the mechanisms underlying these assembly rules remains challenging due to host-mediated confounding factors and functional redundancy among phylogenetically divergent microbes^3^. Bottom-up synthetic consortia, such as the 12-member Oligo-Mouse-Microbiota (OMM12), provide a tractable framework to bypass these hurdles and map defined metabolic interactions^4,5^. While key nutritional niches were identified *in vitro* and *in vivo^6–8^*, direct causal links between *in vitro* metabolic mechanisms and spatial *in vivo* responses during dietary interventions remain lacking.

This gap is particularly critical at the mucosal barrier, where dietary fiber deprivation forces versatile generalists to shift to host mucin foraging, driving mucosal barrier degradation and dysbiosis^9–11^. Although prebiotic fibers are deployed to counteract this thinning and reinforce barrier integrity, therapeutic outcomes remain notoriously variable. Because fiber processing relies on interspecies division of labour and cross-feeding cascades, isolated monoculture screens fail to predict emergent community behaviours^12–15^. Decoding fiber-microbe specificity is essential for building mechanistic degradation models and rationally designing prebiotics to fortify the mucosal barrier^16^. Furthermore, standard *in vitro* cultivation models routinely omit the spatial and nutritional architecture of the mucus layer, limiting their capacity to predict *in vivo* dynamics. To bridge this divide, recent advances have incorporated mucin in an agar-bound form, retaining significantly higher species diversity and successfully mimicking the mucosal community found *in vivo*^17–19^. While these findings underscore the vital triad of mucin, fiber, and microbial interactions, the dynamic interplay of bacteria during prebiotic administration has remained largely correlative rather than causally defined.

To establish a causative, spatially relevant framework for prebiotic processing, we coupled a modified OMM12 consortium with an *in vitro* mucin-carrier platform that mimics the physical mucosal microenvironment. Soluble xylooligosaccharides (XOS) are widely utilized prebiotics known to selectively modulate the gut microbiota, restore mucosal health, and reduce inflammation^20,21^. While numerous studies have reported the enrichment of *Bifidobacteriaceae* upon XOS supplementation *in vitro* and *in vivo*^22–24^, its impact on a complex bacterial consortium remains poorly understood. We integrated 16S metabarcoding, targeted metabolomics, and community metatranscriptomics to map XOS degradation cascades and identify distinct primary utilizers. Validating this platform *in vivo*, gnotobiotic OMM12 mice fed a fiber-free diet with or without XOS replicated the rapid 24-hour compositional equilibrium observed *in vitro*, marked by a specific expansion of *Blautia coccoides*. Crucially, core XOS-degrading gene expression profiles demonstrated strong concordance between *in vitro* cultures and colonic environments. By applying spatially resolved transcriptomics along the proximal-to-distal colonic axis, we disentangled intrinsic substrate preferences from host-mediated microenvironmental imprints shaping bacterial expression. Together, our work provides a scalable, highly predictive platform that that offers a scalable blueprint for testing a wide array of prebiotics both *in vitro* and *in vivo*.

### Temporal dynamics and stability of an *in vitro* gut community

To investigate how prebiotics drive emergent community dynamics beyond individual monoculture capabilities^12,13^, we established a spatially structured and compositionally stable *in vitro* model using a modified OMM12 consortium (OMM12*). To simulate the spatial architecture of the mucosal layer, we incorporated mucin-agar matrices into a cecum-mimicking media^6,7,18^. These carriers provided a distinct niche for specific bacteria without perturbing overall exometabolite profiles (**Figures S1A &B**). Daily passaging confirmed that the community rapidly attained a stable, reproducible equilibrium within 24 hours (**Figures 1A&B, S1C**) that remained stable over seven days (**Figure 1C**).

**Figure 1:**
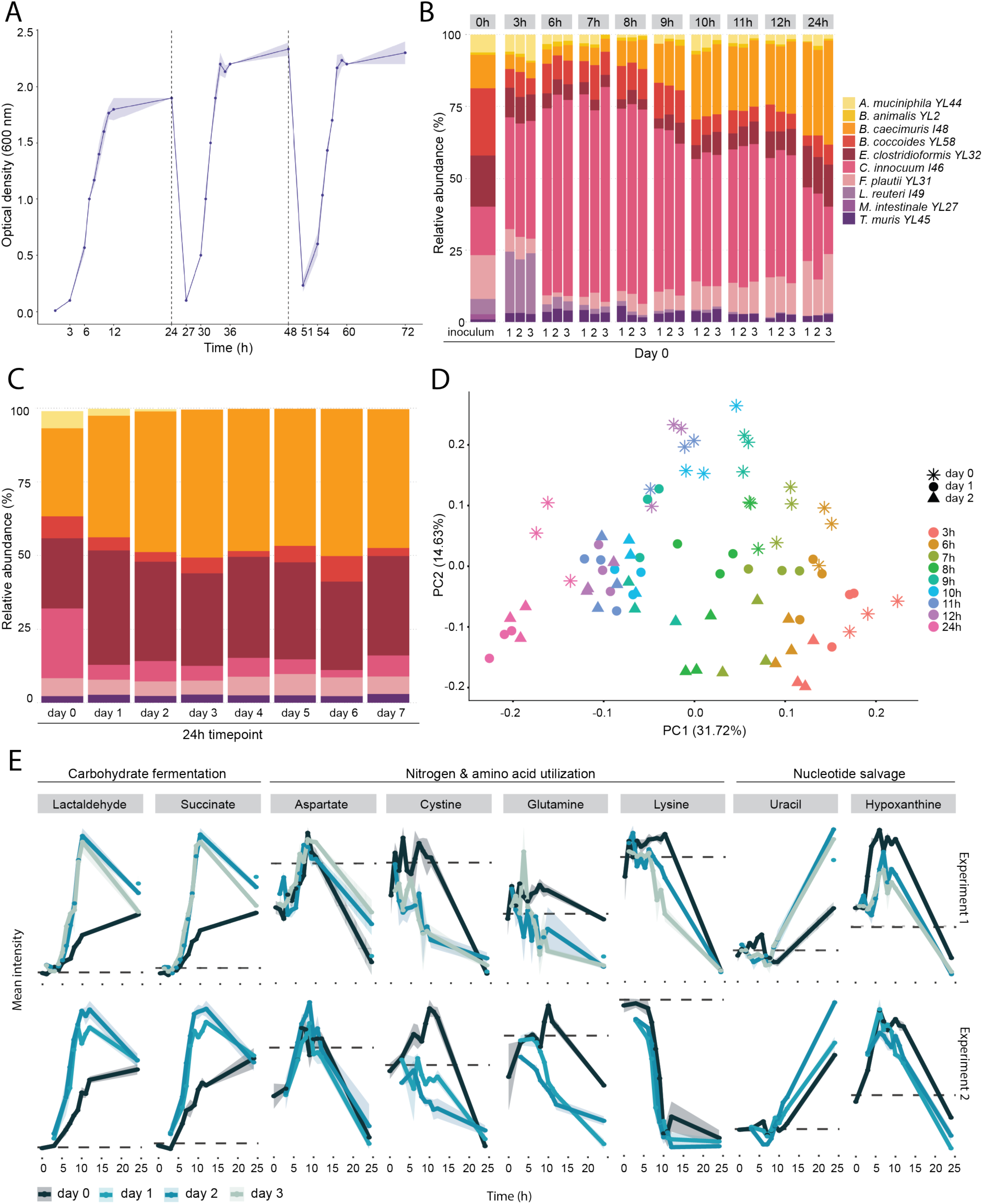
Compositional, functional, and metabolic stabilization of the synthetic OMM12 consortium in a specialized mucin-carrier *in vitro* platform. A) Growth kinetics of the synthetic OMM12 consortium over 24-hour cycles across three consecutive serial passages. Mean values of three replicates with SD are shown. **B)** High-resolution temporal dynamics of relative bacterial abundance based on 16S rRNA gene profiling during the initial 24-hour passage (day 0) in triplicate cultures. **C)** Long-term compositional equilibrium of the OMM12 community at 24-hour harvest time points across seven consecutive daily passages. Mean values of three biological replicates are shown. **D)** Principal Component Analysis (PCA) score plot of untargeted LC-MS/MS metabolomics from culture supernatants tracking metabolic trajectories across sampling time points and consecutive daily passages. Three biological replicates per time point. **E)** Time-course dynamics over three daily passages of selected extracellular metabolites in carbohydrate fermentation, nitrogen and amino acid utilization, and nucleotide salvage. Two independent experiments are shown, each with 3 replicates.

As composition does not necessarily reflect function^25^, we sought to investigate the functional stability of our *in vitro* community. Untargeted metabolomics of culture supernatants confirmed that functional outputs are also reproducibly stable after 24 hours. Principal component analysis (PCA) revealed that day 0 metabolically segregates from the following days along PC2 (14.63%), while PC1 (31.72%) reflects predictable metabolic shifts over time (**Figure 1D**). To validate this functional reproducibility across independent experiments, we tracked the trajectories of eight representative metabolites spanning core physiological pathways. These trajectories were highly consistent across independent experiments, indicating that while day 0 is shaped by community assembly, subsequent days display reproducible metabolic behaviour (**Figure 1E**). Altogether, these data demonstrate a stable *in vitro* community with a robust, and functionally predictable baseline after 24 hours post-inoculation, which can be used to address mechanistic questions regarding how prebiotics shape bacterial communities.

### Addition of XOS leads to the emergence of distinct guilds after 24h and distinct sugar consumption patterns

To determine how a defined substrate shapes community dynamics over time, we tracked growth, composition, and metabolite profiles in our validated *in vitro* platform. Targeted LC-MS of hydrolysed XOS confirmed a carbohydrate backbone composed of 94% xylose and 2% arabinose, with trace galactose (2%) and glucose (2%) likely representing manufacturing artifacts (**Figure 2A**). Although vendor specifications listed methyl-glucuronic acid, its concentration remained below detectable limits (<0.22 µM) despite positive spike-in controls (**Figure S2A**). XOS supplementation to our *in vitro* platform reduced maximal biomass (OD600 ∼1.3 vs. ∼2.3 in control; **Figure 2B**), likely driven by rapid lactate accumulation and subsequent pH stress (**Figures S2B and S2C**). Community composition analysis revealed a distinct structural shift within 24 hours of XOS exposure (**Figure 2C**) that remained stable through 72 hours, in line with our previous observations (**Figure S2D**). Differential enrichment analysis revealed clear condition-specific preferences across the consortium, notably the expected expansion of *Bifidobacterium animalis* (**Figure 2D)** ^23,24^.

**Figure 2:**
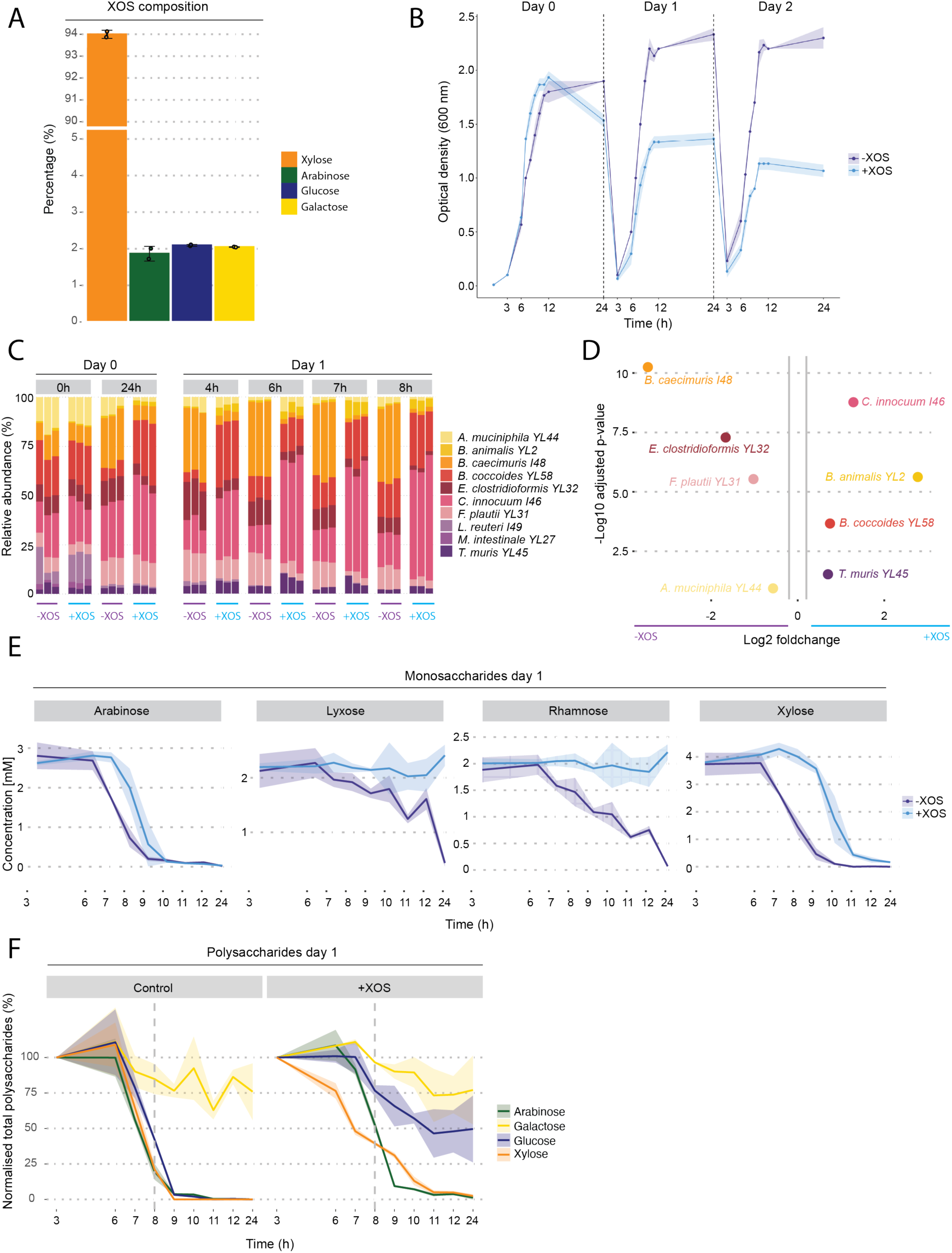
Xylooligosaccharide (XOS) supplementation reorients synthetic consortium composition and carbohydrate fermentation dynamics *in vitro*. **A)** Monosaccharide composition of the XOS substrate. Data are the mean of two technical replicates. **B)** Growth curves of the synthetic OMM12 community across three consecutive passages in APF medium with and without XOS supplementation. Mean values of three replicates with SD are shown. **C)** Dynamic shifts in relative species abundance based on 16S rRNA gene profiling in APF medium with and without XOS supplementation. **D)** Differential abundance of individual OMM12 strains in APF medium with and without XOS supplementation. t-test with Benjamini-Hochberg adjustment; threshold lines indicate |log2FC| > 0.2. **E)** Extracellular concentration of monosaccharides in OMM12 culture supernatants over Day 1. **F)** OMM12 consumption profiles of soluble polysaccharide-derived sugars in APF medium with and without XOS supplementation over Day 1. Dotted line represents the 8h timepoint at which transcriptomic samples were taken. **B,D&F)** Data points are mean values ± SD from three biological replicates.

To map sugar utilization patterns, we quantified both free monosaccharides (**Figure 2E**) and total acid-hydrolyzed sugar fractions (**Figure 2F**). Analysis of unsupplemented media revealed traces of mannose or glucose, likely originating from yeast and soy extracts (**Figure S2E**). The community exhibited highly reproducible sugar degradation across daily cycles (**Figure 2E**). Niche sugars like lyxose and rhamnose were not consumed in the presence of XOS, whereas free arabinose and xylose were rapidly depleted in both conditions, displaying distinct consumption lags under XOS supplementation (**Figure 2E**). Because total carbohydrate concentrations differed substantially between conditions, we normalized each pool to its 3-hour baseline to enable a direct comparison of consumption kinetics across media. By mid-exponential growth (8 hours), the community had consumed ∼50% of the supplemental XOS pool, whereas the minimal background carbohydrate reserves in the control medium were already >75% exhausted (**Figure 2F**). Notably, galactose concentrations, a component of both XOS and mucin, remained steady throughout growth, reflecting a balanced release from the mucin-carriers and slow microbial consumption. Collectively, these kinetics demonstrate that the introduction of soluble prebiotics fundamentally rewires the community, driving shifts in community composition and sugar consumption.

### XOS degradation yields distinct transcriptional responses and substrate uptake strategies

Mapping glycoside hydrolase (GH) families across the OMM12 genomes established distinct genomic potentials for fiber and mucin processing (**Figure 3A**). As expected, *Akkermansia muciniphila* encoded GH families specialized for mucin and galactan processing. Notably, *Bacteroides caecimuris* encoded the broadest CAZyme repertoire, spanning nearly all predicted substrate categories (**Figure 3A**). *Blautia coccoides* also harbored a surprisingly diverse GH profile despite not being traditionally classified as a primary polysaccharide degrader. In contrast, *Clostridium innocuum*, *Flavonifractor plautii*, and *Turicimonas muris* possessed sparse GH profiles, indicating their ecological roles within the community are independent of fiber and mucin cleavage.

**Figure 3:**
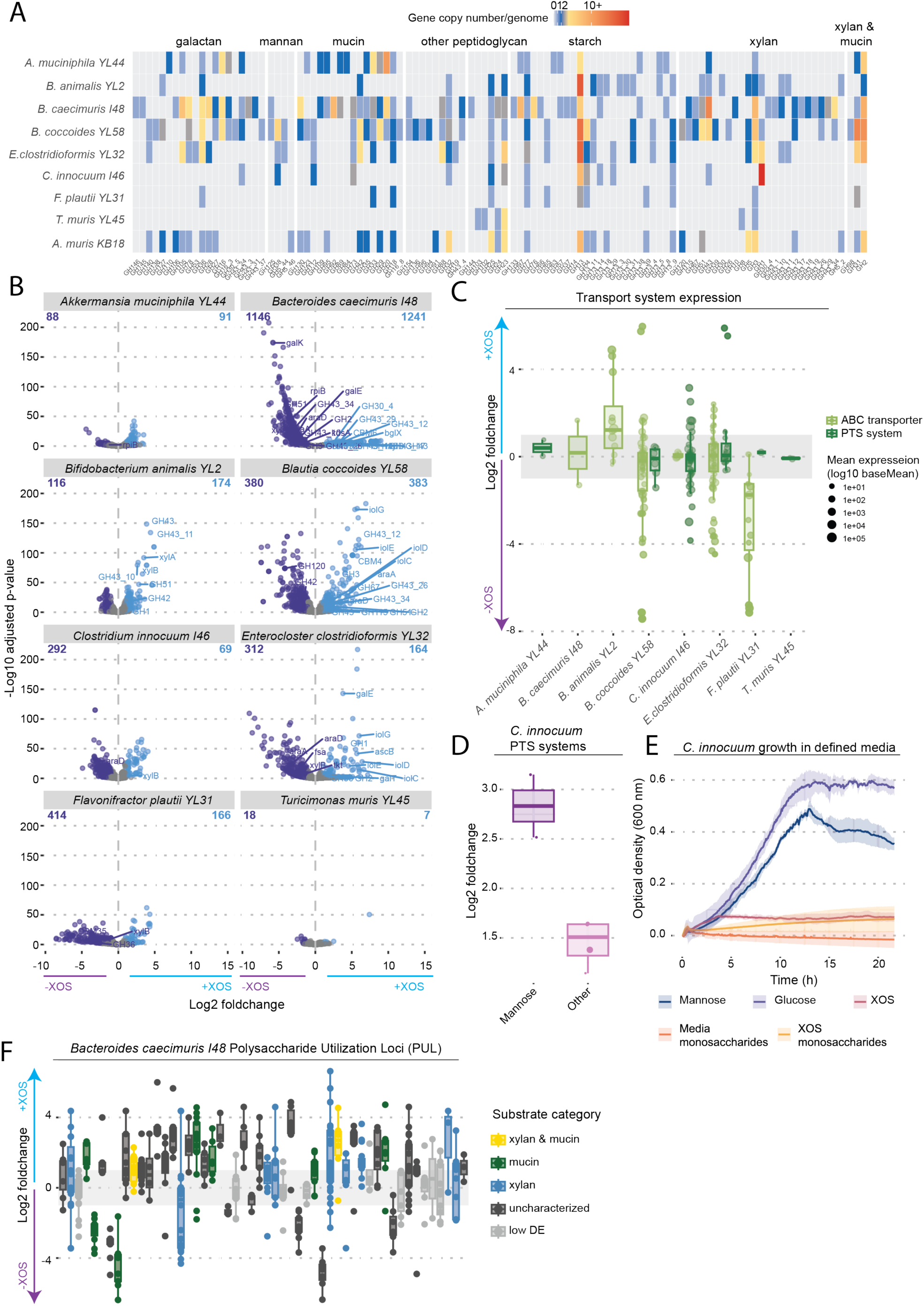
Transcriptional profiling identifies strain-specific glycan transport and utilization machinery in response to XOS. **A)** Genomic distribution heatmap of Carbohydrate-Active Enzyme (CAZyme) copy numbers across OMM12 genomes. **B)** Differential gene expression in individual OMM12 community in APF medium with and without XOS supplementation. Differentially expressed genes (DEGs) defined by adjusted p-values < 0.05 and |log2FC| > 1] from from four biological replicates. Total DEG counts are indicated in each upper quadrant. Genes marked in blue are differentially expressed in media supplemented with XOS whereas genes in purple are DEG in the control**. C)** Expression shifts of carbohydrate transport systems across OMM12 strains. Circle size reflects baseline gene expression level from four biological replicates. **D)** Increased expression of mannose-specific PTS components and other PTS transporters in *C. innocuum* grown community in XOS-supplemented APF medium relative to the control. **E)** Growth profiles of *C. innocuum* in defined media supplemented with different carbon sources. Curves represent mean values ± SD from three biological replicates. **F)** Differential gene expression of Polysaccharide Utilization Loci (PULs) in *Bacteroides caecimuris I48*. They are categorized by target substrate specificity. PULs in the gray zone are not differentially expressed (low DE). Boxplots show median, IQR, and range from four biological replicates.

To resolve individual species contributions to XOS, we profiled mid-exponential (day1) metatranscriptomes, normalizing sequence reads against relative species abundances to reflect genuine transcriptional regulation^26^. This sampling window was selected to capture primary carbohydrate responses while minimizing confounding acid stress. Nonetheless, subtle stress-associated pathways such as molecular chaperones and ATP synthase subunits were already upregulated in five of the eight community members (**Figure S3A**). These early stress responses occurred during active substrate processing rather than carbon exhaustion, as substantial pools of free arabinose (6 mM) and xylose (7 mM) remained in the medium (**Figure S3B**). While some members remained largely unaffected—such as *T. muris* (18 genes downregulated, 7 upregulated)—others underwent extensive transcriptomic rewiring, most notably *Ba. caecimuris*, *Bl. coccoides*, and *Enterocloster clostridioformis* (**Figure 3B**).

Substrate transport architectures mirrored these divergent functional roles. Strains employ two fundamental transport mechanisms: phosphoenolpyruvate (PEP)-dependent phosphotransferase systems (PTS), which maximize bioenergetic efficiency by phosphorylating simple monosaccharides during translocation, and ATP-binding cassette (ABC) transporters, which consume ATP to import intact oligosaccharides against steep concentration gradients^27^. Reflecting active sequestration of intact short-chain oligosaccharides, *Bl. coccoides* and *Bif. animalis* strongly upregulated multi-sugar ABC transporters in response to XOS supplementation (**Figures 3C**)^28^, whereas *F. plautii* downregulated its ABC transport machinery (**Figure 3C**). In contrast, PTS systems were enriched in *E. clostridioformis* and *C. innocuum* (**Figure 3C**). *C. innocuum*, which expanded significantly in the presence of XOS, selectively induced mannose-specific PTS transporters (**Figure 3D**). Defined media assays confirmed that *C. innocuum* grew robustly on glucose and mannose but failed to utilize XOS, XOS-derived or media-derived monosaccharides (**Figure 3E**). Thus, the bloom of *C. innocuum* in XOS-supplemented environments is driven by a mannose-mediated niche expansion rather than direct prebiotic catabolism.

The profound transcriptional response of *Ba. caecimuris* coincides with a severe reduction of its relative abundance in XOS-containing media. In *Bacteroides* species, carbohydrate-active enzymes (CAZymes) are organised in Polysaccharide Utilization Loci (PULs), co-regulated genetic islands specialized in sugar degradation. Profiling PUL expression by predicted substrate specificity revealed a highly coordinated, operon-level activation across multiple loci (**Figure 3F**). While several xylan-targeted loci were significantly induced, *Ba. caecimuris* also strongly upregulated an array of mucin-degrading and uncharacterized PULs (**Figure 3F**). Because the supplied XOS consists of low-molecular-weight oligosaccharides (300-900 Da), this extensive PUL response likely represents an energetically inefficient strategy. Previous studies also confirm that longer-chain XOS preferentially enrich *Bacteroides* species, whereas shorter-chain XOS favour members of the *Bifidobacteriaceae* and *Lachnospiraceae* families that are specialized for the rapid uptake and metabolism of low-molecular-weight carbohydrates^23,24^. Altogether, XOS elicits specialised transcriptional changes across consortium strains,, showcasing how prebiotic additions reshape community structure by directly altering substrate utilization and indirectly enabling niche exploitation.

### Hierarchical processing of XOS

Integrating transcriptomic responses with genomic annotations revealed that only four strains significantly induced XOS degradation and metabolism machinery upon supplementation, namely *Ba. caecimuris*, *Bif. animalis*, *Bl. coccoides*, and *E. clostridioformis* (**Figure 4A**). *Ba. caecimuris* predominantly upregulated extracellular degradation enzymes (xylanases and arabinofuranosidases) rather than internal monosaccharide catabolism (**Figure 4A**). Conversly, *E. clostridioformis* displayed a scavenger profile, exclusively upregulating galactose and inositol utilization pathways without contributing to XOS cleavage (**Figure 4A**). Meanwhile, *Bif. animalis* and *Bl. coccoides* displayed complementary utilization profiles: *Bif. animalis* prioritized xylose catabolism alongside arabinofuranosidases, whereas *Bl. coccoides* upregulated arabinose operons and targeted xylosidases, pointing to a coordinated cross-feeding network (**Figure 4A**).

**Figure 4:**
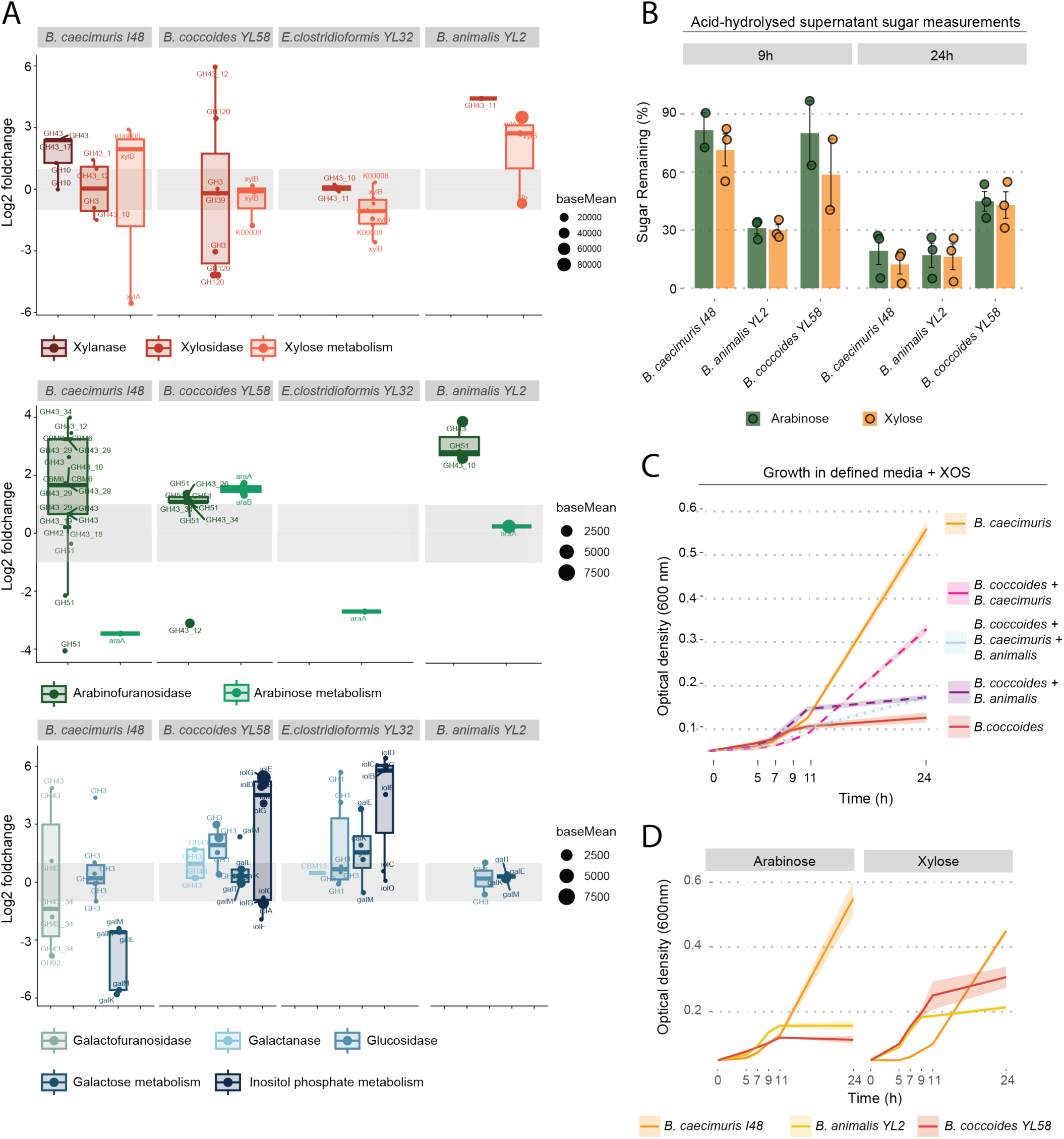
Susbtrate division among primary XOS degraders. **A)** Differential expression of selected catabolic operons in strains responding to XOS supplementation. The operons include xylanases, xylosidases, xylose metabolism, arabinofuranosidases, arabinose metabolism, glucuronidases, glucuronate metabolism, and inositol phosphate pathways.Point size scales with baseMean expression and boxplots indicate median and IQR of four biological replicates. **B)** Relative consumption profiles of acid-hydrolyzed supernatant arabinose (green) and xylose (orange) during growth of primary degraders in defined media with XOS. Data are means ± SD from three technical replicates. **C)** Growth curves of *Blautia coccoides* in monoculture and in combination with *Bifidobacterium animalis* and/or *Bacteroides caecimuris* in defined media containing XOS as the sole carbon source. Data are means ± SD for three technical replicates. **D)** Growth curves of primary degraders in defined media supplemented with either arabinose or xylose as the sole carbon source. Data are means ± SD of three technical replicates.

Phenotypic validation in defined XOS media confirmed that all three predicted primary degraders successfully grew, though they exhibited distinct lag phases that likely reflect their divergent enzyme allocation strategies (**Figure S3C**). Supernatant analysis of these monocultures from exponential (9h) and stationary (24h) phases revealed that *Bif. animalis* rapidly depleted ∼70% of both xylose and arabinose by 9h, whereas *Ba. caecimuris* and *Bl. coccoides* exhibited slower degradation (**Figure 4B**). Notably*, Bl. coccoides* left > 40% of the initial sugars unconsumed at 24h, demonstrating incomplete independent XOS degradation potential (**Figure 4B**). Consistent with this physiological limitation, *Bl. coccoides* achieved a significantly higher maximal biomass (OD600) when cocultured with either *Ba. caecimuris, Bif. animalis*, or both (**Figure 4C**). In contrast, *Ba. caecimuris* attained its highest biomass in monoculture, with the addition of coculture partners progressively decreasing its peak biomass (**Figure 4C**). This confirms that *Ba. caecimuris* performs the primary degradation of XOS, liberating public goods without receiving a reciprocal growth advantage, mirroring its competitive decline in rich medium.

Evaluating monoculture growth on arabinose and xylose unraveled the competitive logic behind community substrate partitioning (**Figure 4D**). *Bl. coccoides* intrinsically preferred xylose over arabinose, achieving a shorter lag phase and higher biomass on xylose (**Figure 4D**). However, its growth trajectory on xylose overlapped with *Bif. animalis*, which outcompetes *Bl. coccoides* on this substrate through its bioenergetically efficient phosphoketolase pathway. This metabolic dominance forces *Bl. coccoides* to upregulate its arabinose utilization operon in community setting despite its preference for xylose. Together, these interactions outline a clear XOS degradation cascade driven by initial breakdown, competition for free xylose and metabolic adaptation.

### *In vivo* supplementation of XOS largely reflects *in vitro* community shifts

To evaluate how our *in vitro* findings translate to a host environment, gnotobiotic OMM12 mice were switched from standard chow to a fiber-free diet with or without 10% XOS supplementation (**Figure 5A, S4A**). Diet switches caused transient, minor weight loss (∼2.5%) that stabilized by day 2 across both groups (**Figure S4B**). Within 24 hours, two distinct, highly stable community structures established and persisted over five days (**Figure 5B**). Fiber deprivation alone induced a pronounced expansion of the mucolytic *A. muciniphila^10^*, whereas XOS supplementation led to massive structural reorganization dominated by *Bl. coccoides* (∼70% relative abundance), reducing overall alpha diversity **(Figure 5B, S4C).**

**Figure 5:**
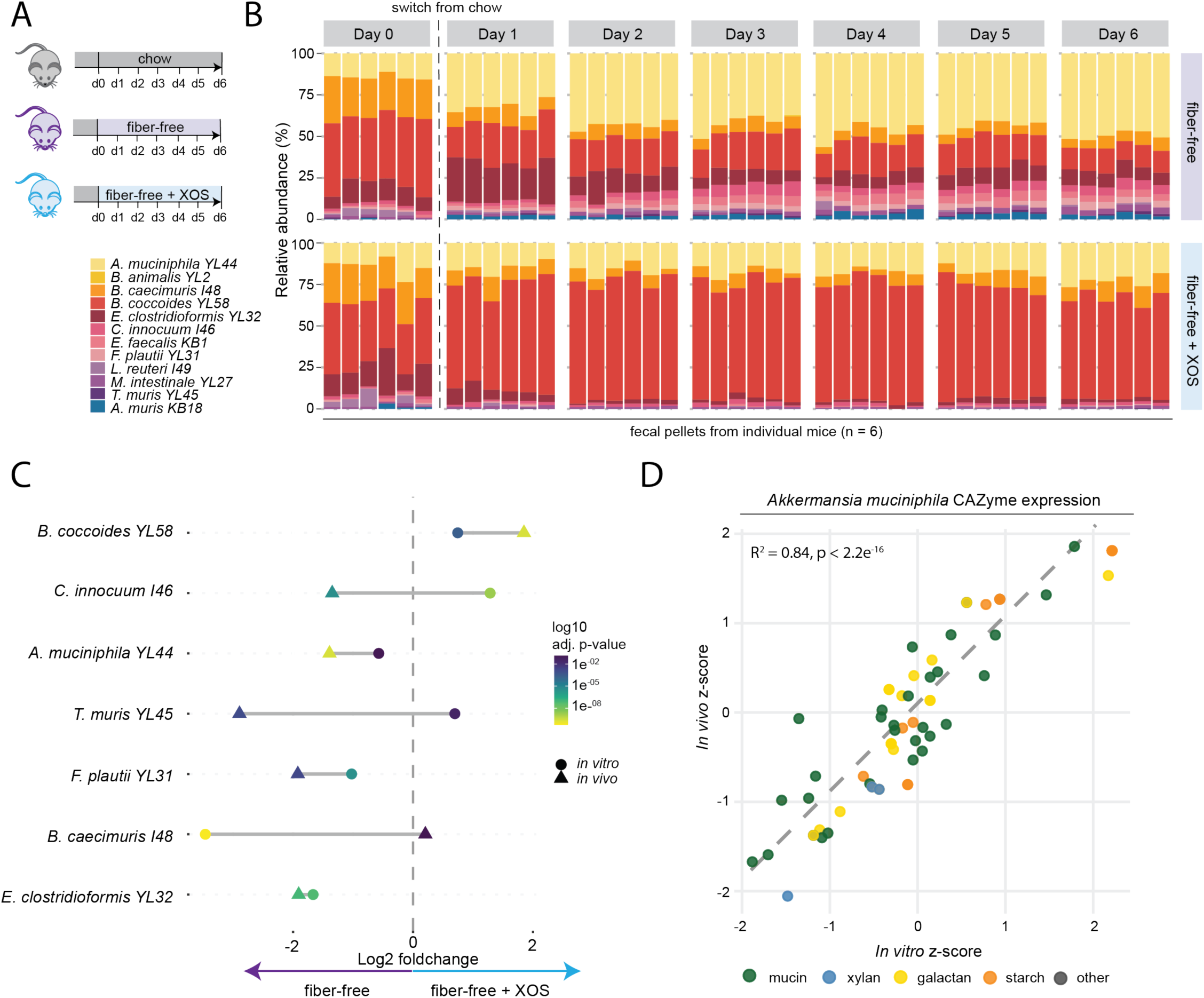
*In vivo* XOS supplementation diverts the OMM12 consortium from mucosal foraging to prebiotic processing. **A)** Schematic representation of the *in vivo* dietary study design using gnotobiotic OMM12 mice (N=6 mice per group) transitioned on Day 0 from standard chow to either a fiber-free diet or a fiber-free diet supplemented with 10% XOS over 6 days. **B)** 16S rRNA-based tracking of the fecal microbiome composition in individual mice. **C)** Species-specific enrichment in *in vitro* mucin-carrier cultures (circles) and *in vivo* fecal samples (triangles) in fiber-free and XOS-supplemented conditions. T-test with BH-correction. **D)** Functional validation of mucosal mimicry. Scatter plot correlating *A. muciniphila* CAZyme gene expression (z-score) *in vitro* (mucin-carrier platform) versus *in vivo* (colonic transcriptomes) across the indicated substrate classes. Pearson correlation, R^2^ = 0.84, p < 2.2e^-16^.

Comparing species enrichment between *in vitro* and *in vivo* settings revealed strong directional concordance for four of seven shared species (*Bl. coccoides*, *A. muciniphila*, *F. plautii*, and *E. clostridioformis*; **Figure 5C**). Notable host-specific deviations included *Bif. animalis*, which remained below detection in adult mice at baseline^6,29^ and *Acutalibacter muris*, which expanded exclusively *in vivo* during fiber deprivation, in line with its mucin-targeted CAZyme repertoire (**Figure 3A, 5B**). Conversely, *Ba. caecimuris, T. muris,* and *C. innocuum* displayed higher relative competitiveness *in vitro*(**Figure 5C**).

To test whether our *in vitro* mucin carriers accurately simulate mucosal microenvironments, we compared normalized *A. muciniphila* CAZyme expression (z-scores) between carrier matrices and *in vivo* colonic transcriptomes (**Figure 5D**). CAZyme profiles across distinct glycan classes correlated strongly between conditions (**Figure 5D**, R² = 0.84). The strong similarities in taxonomic trajectories and mucosal CAZyme expression underscores the utility of our *in vitro* mucin-carrier system for modeling complex gut microbiome dynamics.

### Spatial gradient of prebiotic fermentation highlights intra-host functional plasticity of *B. coccoides*

The chemical architecture of dietary fibers dictates their spatial fermentation dynamics along the gastrointestinal tract ^16^. Given its low molecular weight (300–900 Da) and linear backbone, XOS represents a typical proximally fermented substrate. Because glucose and galactose predominate in the fiber-free diet and host mucosal layer respectively, we focused on quantifying the primary XOS monomers, xylose and arabinose. Targeted quantification of acid-hydrolyzed carbohydrates from cecal (proximal) and fecal (distal) contents confirmed that XOS-derived sugars were significantly enriched in the cecum (**Figures 6A & S5A**), with xylose levels markedly elevated exclusively in XOS-supplemented mice (**Figure 6A**). Spatial community profiling revealed that taxonomic composition remained remarkably uniform along the colonic axis despite the proximal-to-distal sugar gradient (**Figures 6B & S5B**). *E. clostridioformis* exhibited consistent proximal enrichment regardless of diet (**Figure S5B**), reflecting a specialized cecal niche driven by its butanoate metabolic pathway, which couples butyrate conversion to succinate for biomass generation (**Figure S5C**)^30^.

**Figure 6:**
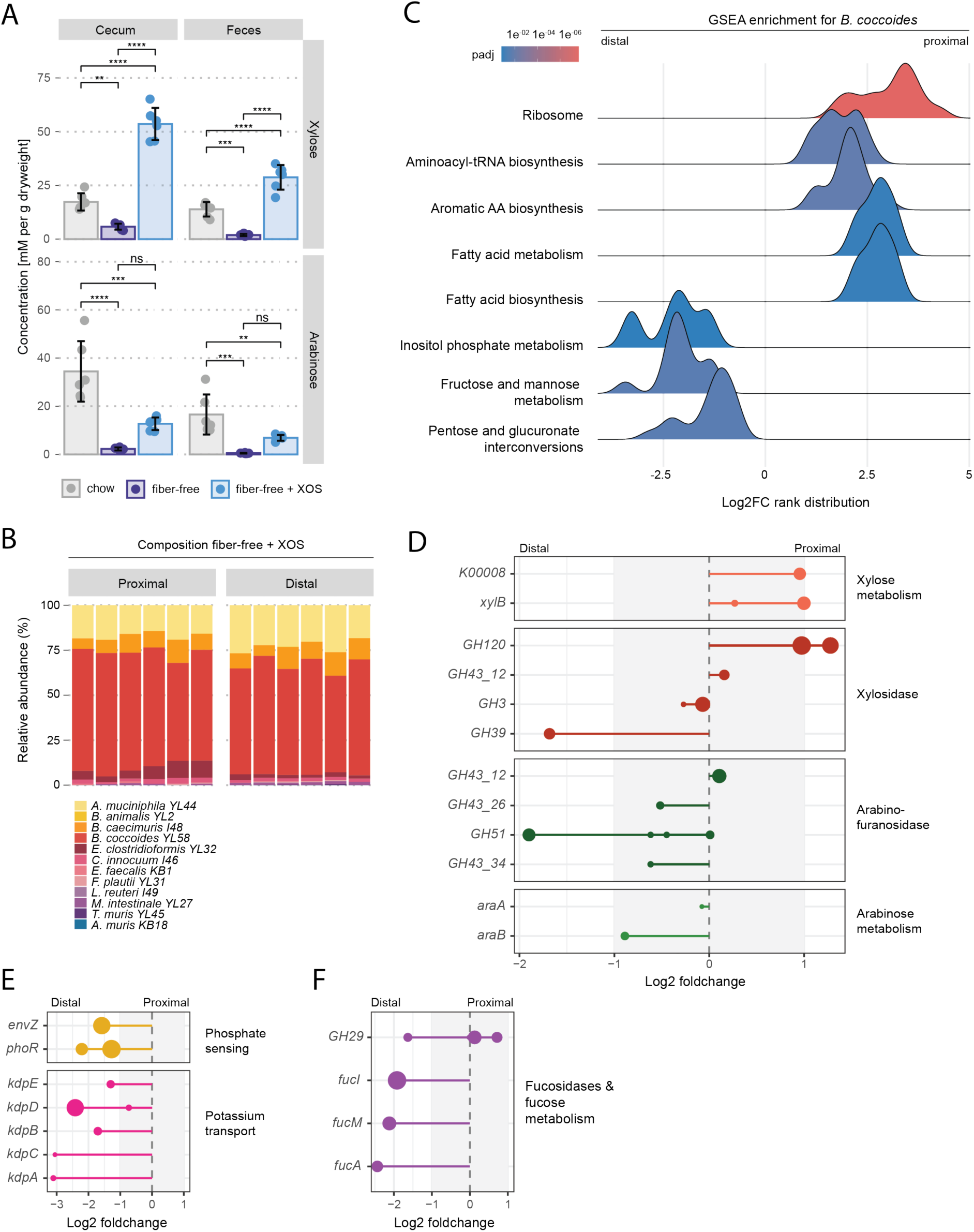
Spatial gradients of XOS depletion drive longitudinal transcriptional and metabolic adapations along the colonic axis. **A)** Acid-hydrolyzed polysaccharide-derived sugar concentrations in cecum (blue) and feces (orange). Data are means ± SD of six mice per group. Global statistical significance was determined by One-Way ANOVA, followed by Tukey’s post-hoc test for pariwise comparisons; **p < 0.01, ***p < 0.001, ****p < 0.0001, ns = not significant. **B)** 16S RNA-based spatial community composition in proximal and distal colonic luminal contents of mice fed a fiber-free or XOS diet. **C)** Gene Set Enrichment Analysis (GSEA) ridge plot depicting functional pathways significantly enriched in *B. coccoides YL58* between the distal and proximal colonic niches. Rank distribution is based on log2 foldchanges and the color gradient indicates adjusted p-value (padj). **D)** Differential spatial expression of *B. coccoides* genes involved in xylose metabolism and arabinose catabolism. Data are means of three mice. **E)** Differential gene expression of *B. coccoides* fucosidases and fucose metabolism operons in the colon. Data are means of three mice. **F)** Differential gene expression of *B. coccoides* inorganic phosphate sensing and potassium transport operons in the colon. Data are means of three mice.

As static taxonomic composition fails to capture localized metabolic activity, we investigated how consortium members adapt to progressive substrate depletion during transit through the lower gut. Paired transcriptomic profiling of matched proximal and distal samples within individual XOS-supplemented mice enabled us to map longitudinal functional shifts while controlling for host-to-host variation, leveraging the ∼2-hour colonic transit providing an adequate physiological window for gene regulatory shifts^31^. Metatranscriptomic profiling revealed that *Bl. coccoides* mounted the most extensive spatial shift, altering 596 genes in the proximal colon and 658 genes in the distal colon (**Figure S5D**). Unbiased Gene Set Enrichment Analysis (GSEA) mapped a distinct functional transition: *Bl. coccoides* prioritized anabolic growth and central metabolic pathways in the proximal colon, but switched to nutrient starvation responses like alternative carbon utilization pathways in the distal colon (**Figure 6C**). Strikingly, these spatial expression profiles directly recapitulated the physiological behaviors established in our *in vitro* assays. The transcriptome of *Bl. coccoides* revealed its intrinsic preference for xylose over arabinose. In the absence of its competitor *Bif. animalis in vivo* it selectively upregulated xylosidases and the xylose utilization operon in the proximal colon before inducing arabinose catabolism in the distal colon (**Figure 6D**). Conversely, *Ba. caecimuris* mirrored its *in vitro* phenotype by inducing an extensive array of arabinofuranosidases and xylanases in the proximal colon without upregulating downstream catabolism pathways (**Figure 4A, S5E**).

As XOS availability is significantly reduced in the distal colon, *Bl. coccoides* encountered severe inorganic phosphate and potassium starvation (**Figure 6E**). This phosphate restriction likely arises from the extensive microbial consumption during the proximal growth surge. To overcome the carbon source exhaustion, *Bl. coccoides* induced its fucose utilization operon in the distal colon, effectively pivoting from XOS fermentation to scavenging host-derived mucin glycans liberated by primary mucin degraders (**Figure 6F**). Together, these data demonstrate high functional predictability between *in vitro* models and *in vivo* dynamics. Furthermore, they highlight the necessity of accounting for longitudinal microenvironments when designing prebiotic therapies: the sequential transition of *Bl. coccoides* from proximal XOS fermentation to distal mucin scavenging underscores a remarkable metabolic plasticity that underpins its competitive dominance across the entire colon.

## Discussion

Dietary fibers prevent microbiota-driven degradation of the mucosal barrier^10^. However, the mechanisms governing fiber-microbe-mucin interactions within complex gut communities remains poorly understood^16^. Here, we establish a stable, reductionist *in vitro* community using a mucin-carrier platform to resolve interspecies prebiotic processing and mucosal protection. The community achieves compositional and functional equilibrium within 24 hours, mirroring the rapid stabilization observed *in vivo* following dietary transitions (**Figure 1B, S1C, 5B**). The high correlation in *A. muciniphila* CAZyme gene expression between *in vitro* mucin carriers and *in vivo* colonic transcriptomes (R² = 0.84; **Figure 5D**) validates these carriers as accurate physiological mimickry for the host mucosal niche. Extending our multi-omic approach along the colonic axis *in vivo* revealed how spatial XOS gradients drive profound longitudinal metatranscriptomic shifts, capturing both core XOS degradation machinery and dynamic bacterial adaptations to regional host microenvironments (**Figure 4A, 6D-F, S5C-E**). Consequently, this validated system enables strain-resolved mechanistic studies of fiber processing while capturing critical host-microbe interactions.

Integrating 16S rRNA profiling, targeted metabolomics, and species-normalized metatranscriptomics illustrates how fiber chemical architecture dictates microbial competition. *Blautia producta* has previously been shown to be able to grow on XOS using xylosidases^32^, which is reflected in our community by *Bl. coccoides* both *in vivo* and *in vitro* (**Figure 4A, S3C, 5B, 6D**). *Ba. caecimuris* upregulates PULs, xylanases, and mucin-degrading machinery, but not internal monosaccharide catabolism, leading to its competitive decline (**Figure 3F, 4A, S5E**). Conversely, both *Bif. animalis* and *Bl. coccoides* profit from *Ba. caecimuris* degradation (**Figure 3C, 4A&C**).

Substrate competition further shapes community structure across distinct environments. *In vitro*, *Bif. animalis* efficiently metabolizes xylose via the phosphoketolase pathway, competing directly with *Bl. coccoides* (**Figure 4D**), whereas *in vivo*, where *Bif. animalis* is absent, *Bl. coccoides* monopolizes free xylose and dominates the community (**Figure 5B**). While *Bl. coccoides* monocultures cannot fully deplete XOS, coculture with *Ba. caecimuris* enhances overall XOS consumption and biomass production, reflecting an exploitative cross-feeding dynamic (**Figure 4B&C**). This cooperative dependency explains why *Ba. caecimuris* is maintained within the community both *in vitro* and *in vivo*. Although *Bifidobacterium* expansion is a hallmark of XOS supplementation and widely recognized for its therapeutic short-chain fatty acid production^22,33,34^, its natural absence in adult OMM12 mice preempted us from validating this *in vitro* expansion *in vivo*^29^

Dietary fiber deprivation forces the microbiota into host mucosal foraging, marked by *A. muciniphila* expansion and distal short-chain fatty acid depletion (**Figure 5B, S4D**). Paradoxically, the fiber-free diet significantly increased alpha diversity compared to standard chow (**Figure S4C**), highlighting that community diversity alone is an unreliable proxy for gut health^35^. XOS supplementation reversed these pathological shifts partly by fully restoring fecal acetate and propionate (**Figure S4D**). The expansion of *Bl. coccoides* carries significant physiological implications because its acetate and propionate production stimulates mucosal layer production through the activation of the host free fatty acid receptor 2 (*Ffar2*)^36^. *In vitro* fucose supplementation fuels propionate synthesis by *Bl. coccoides*, aligning with its selective upregulation of fucose utilization operons in the distal colon and suggesting a shared metabolic axis *in vivo* (**Figure 6F**). Beyond short-chain fatty acid generation, the conversion of toxic long-chain unsaturated fatty acids into hydroxy-fatty acids by *Blautia* exerts key immunomodulatory benefits on the intestinal epithelium^37^. Promoting *Bl. coccoides* via targeted prebiotics like XOS thus offers a compelling strategy over single-strain probiotic therapies to reinforce intestinal homeostasis.

While the spatial taxonomic composition remains largely uniform along the colon, aside from proximal *E. clostridioformis* enrichment, metatranscriptomics uncovers profound longitudinal functional plasticity (**Figure 6B, S5B & D**). As XOS is progressively depleted during transit, *Bl. coccoides* undergoes a sequential metabolic transition, prioritizing anabolic growth and rapid XOS fermentation via xylosidase and xylose operon induction in the proximal colon, while pivoting to arabinose catabolism and host fucose scavenging in the distal colon (**Figure 6A & B, D&F**).

Altogether, this study establishes a translational platform that bridges reductionist *in vitro* modeling with complex *in vivo* mucosal dynamics. By resolving how fiber architecture and regional microenvironments dictate interspecies interactions to prevent host mucosal foraging, our work provides a mechanistic blueprint for prebiotic processing along the colonic axis. Ultimately, accounting for these longitudinal metabolic cascades will be essential for designing targeted microbiome therapeutics that fortify intestinal barrier function and preserve gut homeostasis.

## Methods

### Bacterial strains and growth

#### Rich Media Preparation

All community experiments were conducted using a modified APF medium adapted from ^7^ (**Table S23**). The medium consisted of 18.5 g/L brain heart infusion (glucose-free), 15 g/L tryptone soy broth (glucose-free), 5 g/L yeast extract, 0.62 g/L K2HPO4, 1 mg/L haemin, 2.5 g/L of a sugar mixture (composed of arabinose, lyxose, rhamnose, and xylose in a 1:1:1:1 ratio), 0.5 mg/L menadione, 3% (v/v) heat-inactivated fetal bovine serum, 0.5 g/L cysteine-HCl‧H2O, and 0.34 g/L NaH2PO4 in ddH2O. For experiments investigating fiber degradation, the modified medium was supplemented with 2 g/L corncob xylan (Biosynth, YC05381).

#### Defined media preparation

Monoculture and coculture validation experiments were performed in a chemically defined medium containing vitamins, minerals, amino acids, and nucleotides (composition detailed in **Table S24**). To assess substrate-specific growth kinetics and substrate preferences, the basal defined medium was supplemented with carbon sources at a final concentration of 5g/L.

Tested carbon sources included individual carbohydrates—xylooligosaccharides (XOS), glucose, mannose, pure arabinose, or pure xylose—as well as defined monosaccharide combinations, including a XOS-derived monosaccharide mixture (xylose, arabinose, and galactose) and a media monosaccharides (arabinose, xylose, rhamnose, and lyxose). All media were filter-sterilized (0.22uM) and reduced to anaerobic conditions prior to inoculation.

#### Preparation of Spatial Mucin Carriers

To simulate the mucosal layer, spatial elements containing mucin-agar gels (adapted from ^18^) were incorporated into all experiments. Briefly, a solution of 0.5% (w/v) mucin (Sigma Aldrich, M2378) and 0.8% (w/v) agar was prepared in sterile ddH2O. To preserve the integrity of mucin ^38^, sterilization was achieved by bringing the mixture to a boil three times in a microwave, with 5-minute cooldown periods between each boiling cycle ^39^. The sterilized mixture was then poured into Petri dishes containing K1 filter media (Koyzentrale, K1-50L) to coat the filters. Both the liquid media and the mucin carriers were reduced in an anaerobic chamber (Coy Laboratory Products) for 24 hours prior to assembling the experimental hungate tubes and bottles. Mucin carriers were added to the hungate bottles at a concentration of 1 carrier per mL of culture volume.

#### Bacterial Cultivation and Inoculum Preparation

All cultivation and experimentation were performed under strictly anaerobic conditions (5% H2, 5% CO2, 90% N2). Bacterial strains (**Table 1**) were recovered from frozen stocks, inoculated into hungate tubes containing modified APF medium, and grown as monocultures for 24 hours at 37°C with shaking at 100 rpm. To prepare the community inoculum, monocultures were adjusted to an optical density (OD600) of 0.1 and mixed in equal volumetric ratios. To optimize long-term community stability, we excluded *Enterococcus faecalis* KB1, which tends to dominate batch cultures despite its low abundance *in vivo*^40^, as well as *Acutalibacter m*uris KB18, which failed to reach detectable growth in our validation trials.

**Table 1:** Bacterial strains.

| **Strain name** | **Abbreviation** | **DSM-Nr.** | **Phyla** | **Gram staining** |
|---|---|---|---|---|
| *Bacteroides (Ba)caecimuris* | I48 | 26085 | Bacteroida | Gram-negative |
| *Muribaculum intestinale* | YL27 | 28989 | Bacteroida | Gram-negative |
| *Akkermansia muciniphila* | YL44 | 26127 | Verrucomicrobia | Gram-negative |
| *Bifidobacterium (Bif) animalis* | YL2 | 26074 | Actinobacteria | Gram-positive |
| *Turicimonas muris* | YL45 | 26109 | Proteobacteria | Gram-negative |
| *Flavonifractor plautii* | YL31 | 26117 | Firmicutes | Gram-positive |
| *Acutalibacter muris* | KB18 | 26090 | Firmicutes | Gram-positive |
| *Enterocloster clostridioformis* | YL32 | 26114 | Firmicutes | Gram-positive |
| *Blautia (Bl) coccoides* | YL58 | 26115 | Firmicutes | Gram-positive |
| *Clostridium innocuum* | I46 | 26113 | Firmicutes | Gram-positive |
| *Enterococcus faecalis* | KB1 | 32036 | Firmicutes | Gram-positive |
| *Limolactobacillus reuteri* | I49 | 32035 | Firmicutes | Gram-positive |

#### Experimental Design and Sampling

Experiments were conducted in biological triplicate. Hungate bottles containing 50mL media and mucin carriers were inoculated with the mixed community to a starting OD600 of 0.01. Cultures were incubated at 37°C with shaking at 100 rpm. Every 24 hours, the liquid part of the community was passaged by transferring a 1:100 dilution of the culture into a new hungate bottle containing fresh media and fresh mucin carriers.

At designated time points, 1 mL samples were drawn from the vessels using a syringe. 100uL of the sample was used for immediate OD600 measurement, and the remainder was centrifuged for 1 minute at 21000 x g. The supernatant was collected and stored for subsequent metabolomic and pH analyses, while the cell pellet was stored at -20°C until DNA extraction. pH measurements were obtained using a refillable glass double-junction electrode (Mettler-Toledo).

### 16S sequencing

#### Sample Collection and DNA Extraction

Bacterial biomass was adjusted to an optical density OD600 of 0.2 in a 100 µL volume. DNA extraction was performed using the Agencourt DNAdvance kit (Beckman Coulter, A48705) according to the manufacturer’s instructions. Briefly, cells were lysed overnight at 55°C with shaking in a lysis buffer containing Proteinase K (NEB, P8107S) and dithiothreitol (DTT). The 100 µL of the lysate was transferred to a new plate and treated with 50 µL of pre-bind buffer. Magnetic beads (Bind-BBE) were added to each sample, incubated for 5 minutes on a magnetic stand, and washed twice with freshly prepared 70% ethanol. The beads were resuspended in 30 µL of elution buffer and incubated for 30 minutes at room temperature before transferring the eluate to a new plate. Extracted DNA was quantified using the Qubit dsDNA BR Assay Kit (Thermo Fisher Scientific) and normalized to 3.33 ng/µL.

#### 16S rRNA Gene Amplification

For amplification of the full-length 16S rRNA gene fragment (V1-V9), 15 ng of DNA was used per reaction and adjusted to a volume of 9 µL with nuclease-free water. Custom primers to ensure amplification of *Bifidobacteriaceae* (Table 2) were added in a 6 µL volume (stock concentration 20 µM each) ^41^. LongAmp Hot Start Taq 2X Master Mix (NEB, M0533S) was added to each well, and a low-cycle PCR was performed (Table 3). PCR products were purified using 0.9x AMPure XP beads (Beckman Coulter, A63881) and eluted in 14 µL of water. Following a second quantification with the Qubit dsDNA BR Assay Kit, DNA concentrations were adjusted to 19.7 ng/µL to provide 178 ng of PCR product for subsequent library preparation.

**Table 2:**
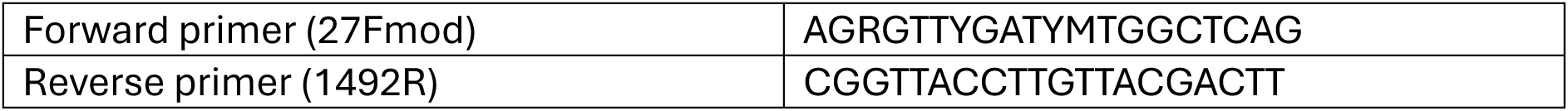
16S primers used.

**Table 3:** PCR program: 95°C 1min, (95°C 20s, 55°C 30s, 65°C 2min) x25, 65°C 5min

#### Library Preparation and Sequencing

Sequencing libraries were prepared using the Native Barcoding Kit 96 (Oxford Nanopore Technologies, SQK-NBD114-96), following the manufacturer’s instructions. The final multiplexed library was loaded onto a MinION Flongle flow cell (FLO-FLG114) and sequenced in-house.

#### Bioinformatics and Taxonomic Profiling

Raw reads generated by MinKNOW (24.06.5) were base called with the high-accuracy model v4.3.0 (400bps). Sequences passing the MinKNOW quality threshold of 9 were concatenated by barcode and filtered using cutadapt (version 5.2) to retain lengths between 1300 and 1700 bp, corresponding to the expected size of the 16S fragment. Filtered reads were mapped against the Emu database (Curry et al., 2022), which contains full-length bacterial 16S sequences. Emu (emu v3.6.2) was executed using 106 threads and a minimum abundance threshold (min_abundance) of 0.00001. Processed outputs and metadata were imported into RStudio (version 4.6.1) for downstream statistical analysis and visualization.

### Transcriptomics

#### DNA and RNA Extraction

Prior to extraction, cell pellets were stored at -80°C to preserve RNA integrity. Simultaneous extraction of genomic DNA and total RNA was performed using the AllPrep PowerFecal Pro DNA/RNA Kit (Qiagen, 80254) with slight modifications. Unless otherwise stated, all centrifugation steps were carried out at 15,000 x g for 1min at room temperature.

Pellets were thawed and resuspended in a mixture of 650 µL CD1 buffer and 100 µL phenol:chloroform:isoamyl alcohol (25:24:1), supplemented with 5 µL of RNase inhibitor (Thermo Fisher Scientific, AM2694). Mechanical lysis was performed using a Precellys homogenizer for 3 cycles of 40 seconds at 6 m/s at 4°C, with 5-minute resting intervals between cycles. Following lysis, the tubes were centrifuged, and the supernatant was transferred to a clean tube. To precipitate proteins, 200 µL of CD2 buffer was added to the lysate, followed by vortexing and centrifugation.

The resulting supernatant was mixed with CD3 buffer in a 1:1 ratio. This mixture was loaded onto an MB DNA Spin Column and centrifuged until the entire volume had passed through, effectively binding the DNA to the membrane. The flow-through, containing the RNA, was collected and mixed with 0.5 volumes of 100% ethanol. This RNA-ethanol mixture was then loaded onto an MB RNA Spin Column and centrifuged.

Both the DNA and RNA columns were subsequently washed with EA buffer and C5 buffer, following the manufacturer’s protocol. A final dry centrifugation step was performed for both columns for 2min at full speed to ensure the complete removal of residual wash buffers. DNA was eluted in 100 µL of nuclease-free water, and RNA was eluted in 50 µL of nuclease-free water.

#### RNA Cleanup, Quality Control, and Sequencing

The eluted RNA was treated with DNase I (Thermo Fisher Scientific, AM2238) for 30min at 37°C to remove residual genomic DNA and subsequently purified using the Monarch RNA Cleanup Kit (50 µg) (New England Biolabs, T2040L). Purified RNA was eluted in 25 µL of nuclease-free water. RNA quantity and integrity were assessed using an Agilent TapeStation system equipped with an RNA ScreenTape (Agilent Technologies) and only samples with a RIN> 7 were considered for sequencing.

Total RNA was shipped to Microsynth AG (Balgach, Switzerland) for library preparation and sequencing. Ribosomal RNA was depleted using the Illumina Ribo-Zero Plus Kit. Sequencing was performed by Microsynth on an Illumina NovaSeq 6000 platform at a targeted depth of 40 million 150bp paired end reads per sample.

### Bioinformatics

#### RNA-Seq Data Pre-processing and Quality Control

Raw RNA sequencing reads were processed using an adapted version of the metatranscriptomics pipeline developed by the Parkinson Lab (Taj et al., 2023),modified for paired-end data. Initial quality control and adapter trimming were performed using Trimmomatic (version 0.39). Illumina Nextera adapters were removed, and reads were quality-filtered using a sliding window of 4:15 and a minimum length cutoff of 50 bp. Vector contamination was subsequently identified and removed by aligning reads to the UniVec database using BWA-MEM (version 0.7.17)

#### rRNA Depletion and Read Alignment

To deplete ribosomal RNA in silico, a custom rRNA database was constructed from the OMM12 strain genomes ^43^. MAFFT (version v7.526) alignments were generated, and covariance models were built using Infernal (version 1.1.5). Paired-end reads were aligned against this custom database, and non-rRNA (putative mRNA) reads were extracted into separate FASTQ files. A pseudogenome was assembled by concatenating the reference genomes of the OMM12 consortium and indexed using BWA. The enriched mRNA reads were mapped to this pseudo genome with BWA-MEM.

#### Read Quantification and Normalization

The resulting alignments were converted to sorted BAM format and quantified using featureCounts (version 2.0.6). Counting was executed in paired-end mode, targeting gene features and aggregating counts by the locus_tag attribute. Sample-specific count matrices were subsequently merged, and genes with zero counts across all samples were filtered out. To account for variations in community composition across samples, raw transcript counts were normalized to the relative abundance of each respective strain within each sample, measured by 16S abundance ^26^.

#### Differential Expression and Functional Annotation

Differential gene expression analysis between the two conditions was performed on the normalized count matrices using the R package DESeq2 (version 1.48.2). Genes with an adjusted false discovery rate below 0.01 and a log2 fold change above 1 were considered as differentially expressed. To facilitate functional annotation, protein-coding sequences (CDS translations) were extracted directly from the GenBank format (.gbff) files for each bacterial strain. KEGG Orthology (KO) assignments were performed using the KEGG Automatic Annotation Server (KAAS) via the bi-directional best hit (BBH) method ^44^ alongside eggNOG-mapper (v3.0.0-beta6) ^45–47^ and koala (version 2026-08-02) ^48^. To specifically profile glycan degradation potential, Carbohydrate-Active enZymes (CAZymes) were annotated using both the dbCAN3 meta-server^49^ and Cayman (v0.13.0)^50^. KEGG Orthology (KO) identifiers were assigned using a majority-consensus strategy across annotations from KofamKOALA, eggNOG-mapper v3 (eggNOG v7.0 database), and KAAS. In the absence of a majority agreement, assignments were hierarchically prioritized based on profile HMM specificity and database currency (KofamKOALA > eggNOG-mapperv3 > KAAS). To resolve ambiguous enzyme activities within GH families, we cross-referenced our CAZyme annotations with BLAST protein domain homologies. Finally, all functional annotations were integrated with the differential expression output tables.

### Metabolomics

#### Untargeted mixed-mode LC-MS

Supernatants were harvested by centrifuging 1mL of culture or extracted sample at maximum speed (21’300g for 1min) and transferring the clarified supernatant to a fresh microplate. Following initial dilution testing to determine optimal analyte coverage within the linear dynamic range, experimental samples were diluted 1:5 (v/v) in LC-MS-grade water. A pooled quality control (QC) study sample was generated by combining equal volume aliquots from each individual sample. Chromatographic separation and mass spectrometry were performed on a ZenoTOF 7600 mass spectrometer (SCIEX) using an Atlantis Premier BEH C18 AX column (1.7uM particle size, 2.1x 30mm; Waters, part no. 186009365). All LC-MS/MS acquisition parameters, mobile phase gradients, and data collection settings were executed as described previously ^51^. Raw spectra were centroided, peak-picked and aligned using masster^52^. The features were annotated on MS1 level using a *Escherichia coli* in-house library with matching retention times and m/z.

#### Untargeted ffow-injection analysis (FIA)

Supernatant samples were diluted 1/100 in MS grade ddH2O prior to injection. An Agilent 6550 QTOF mass spectrometer (Agilent) was used with the method described in Fuhrer *et al^53^*. Samples were run in two batches and injected in a randomized sequence using a Gerstel MPS 3 autosampler (Gerstel). The instrument was run at 4 GHz in negative ionization MS1 mode collecting high resolution mass spectra between 50 and 1000 mass/charge (m/z) ratio. The mobile phase was 60:40 isopropanol:water adjusted to pH 9 using 4 mM NH4F and supplemented with 10 nM hexakis(1H, 1H, 3H-tetrafluoropropoxy)phosphazine and 80 nM taurochloric acid for online mass calibration. Spectra from the two batches (tissue and content) sample runs were aligned and centroided using superalign in matlab. Ions were annotated by matching their measured mass with reference compounds derived from the Human Metabolome Database (HMDB 4.0) subselected for endogenous compounds, fecal and bacterial metabolites and allowing a tolerance of 1 mDa. The resulting data matrix contained 916 ions that could be matched to the deprotonated metabolites from the database. All m/z peaks that remained unmatched were discarded. Ions were further TIC-normalized.

#### Targeted sugar LC-MS

##### Sample Preparation and Sugar Derivatization with PMP

To determine total monosaccharide composition via polysaccharide cleavage, the targeted sugar method described in Sichert *et al* was applied^54^. Briefly, samples were subjected to acid hydrolysis. Briefly, a hydrolysis reagent comprising 2M HCl containing a 1:1000 dilution of an isotopic internal standard mixture (13C-Mannose, 13C-Glucose, 13C-Galactose) was mixed 1:1 (v/v) with 50uL of sample (either undiluted for low-concentration matrices or pre-diluted 1:10 in LC-MS-grade water). To cleave glycosidic bonds, reaction mixtures were incubated overnight at 95°C in a sealed metal clamp block. Following acid hydrolysis, samples were neutralized with28uL 4M NaOH, and 25uL of the neutralized mixture was transferred to a fresh microplate. For free monosaccharide analysis, samples were treated with equivalent volumes of the HCl internal standard solution and NaOH neutralizing solution to maintain matrix consistency, but were processed directly to omit the high-temperature hydrolysis step. Analytical calibration standards were processed identically to match the corresponding sample treatment conditions (hydrolyzed vs. unhydrolyzed).

Monosaccharides were subsequently derivatized by adding 75uL of a freshly prepared derivatization mixture (6.66mL LC-MS-grade methanol, 3.33mL LC-MS-grade water, 174mg 1-phenyl-3-methyl-5-pyrazolone (PMP), and 142uL of 28% NH4OH). The reaction mixture was incubated at 70°C for 100min on a thermal cycler (Bio-Rad), and diluted 1:50 into LC-MS-grade water containing 1% (v/v) formic acid prior to LC-MS analysis. Peaks were identified using Skyline, using the product m/z and retention times outlined in Table 4 & 5.

**Table 4:** external standards.

| **Monosaccharide** | **Product m/z** | **Retention time [min]** | **Concentration highest std [uM]** |
|---|---|---|---|
| Lyxose | 150 | 2.9 | 404 |
| Arabinose | 150 | 5.7 | 505 |
| Xylose | 150 | 5.8 | 505 |
| Rhamnose | 164 | 3.5 | 404 |
| Fucose | 164 | 6.5 | 404 |
| Mannose | 180 | 2.4 | 505 |
| Glucose | 180 | 5.1 | 505 |
| Galactose | 180 | 5.3 | 404 |
| Mannuronic acid | 192 | 2.5 | 74 |
| Glucuronic acid | 192 | 4.9 | 404 |
| Methyl-glucuronic acid | 208 | 5.6 | 500 |

**Table 5:** internal standards.

| **Monosaccharide** | **Product m/z** | **Retention time [min]** |
|---|---|---|
| 13C-Mannose | 186 | 2.4 |
| 13C-Glucose | 186 | 5.1 |
| 13C-Galactose | 186 | 5.3 |

#### LC-MS/MS analysis

Chromatographic separation was performed through gradient elution (Table 6) with the mobile phase made up of two buffers, Buffer A and Buffer B at a flow rate of 1 mL min-1. Buffer A contains LC-MS grade water supplemented with 0.634g/L ammonium formate (Sigma Aldrich, 70221-100G-F) and sonicated for 10 minutes prior to adding 0.1% formic acid (Fisher Scientific, 10596814) and 0.05% medronic acid (InfinityLab Deactivator Additive, Agilent, 5191-3940). Buffer B contains LC-MS grade acetonitrile supplemented with 0.1% formic acid. The first 1.5min of the flow is directed to the waste.

**Table 6:** gradient setup.

| **Total Time (min** | **Flow Rate (uL/min)** | **Buffer A (%)** | **Buffer B (%)** |
|---|---|---|---|
| 2 | 500 | 85 | 15 |
| 7.5 | 500 | 80 | 20 |
| 7.6 | 500 | 0 | 100 |
| 8.5 | 500 | 0 | 100 |
| 8.6 | 500 | 85 | 15 |
| 9.5 | 500 | 85 | 15 |

The ESI source was configured with the following settings: temperature 625°C, a curtain gas level set to 30 (arbitrary units), collision gas to medium, an ion spray voltage of 5500, and Ion Source Gas 1 and Ion Source Gas 2 both set to 90. PMP-derivatives were measured by multiple reaction monitoring (MRM) with previously optimized transitions and collision energies.

#### Targeted SCFA LC-MS

##### Sample Preparation and SCFA Derivatization with Aniline adapted from Chan *et al^55^*

Culture supernatants were diluted 1:5 (v/v) in MS-grade water and subsequently acidified by mixing with 7.5 mM HCl at a 1:2 ratio. To quantify short-chain fatty acids (SCFAs), a standard curve was generated by serially diluting a 1 mM standard mixture (Table 7) in MS-grade water using 1:3 dilution steps down to a final concentration of 0.45 μM. For derivatization, 25 µL of the acidified sample or standard was transferred to a 96-well plate, and 75 µL of freshly prepared derivatization mix (5mL LC-MS grade Methanol, 5mL LC-MS grade water, 91 µL Anilin (Acros Organics, Belgium), 95.5mg of N-(3-Dimethylaminopropyl)-N′-ethylcarbodiimide hydrochloride (EDC, Sigma-Aldrich), 200 µL 2M HCl and 2 µL of internal standard (Table 8)) was added to each well. The derivatization reaction proceeded for 30 minutes at room temperature. The reaction was then quenched by the addition of 10 µL of a β-mercaptoethanol solution (3.5% v/v in MS-grade water).

**Table 7:** External standards (1mM concentration each)

| **Molecule** | **Product m/z** | **Retention time [min]** |
|---|---|---|
| Acetate | 94 | 1 |
| Lactate | 120.3 | 1.1 |
| Propionate | 94.1 | 1.6 |
| Butyrate | 94 | 2.3 |
| Succinate | 176.3 | 2.8 |
| Malate | 174 | 2.7 |
| Fumarate | 94 | 2.5 |

**Table 8:** Internal standards.

| **Molecule** | **Product m/z** | **Retention time [min]** |
|---|---|---|
| Formate_heavy | 94 | 0.9 |
| Acetate_heavy | 94 | 1 |
| Butyrate_heavy | 95 | 2.3 |
| Succinate_heavy | 95 | 2.8 |

##### LC-MS/MS Analysis

Following quenching, the derivatized plates were diluted 1:50 in ddH2O prior to injection. Chromatographic separation and mass spectrometry were performed using an Agilent 1290 Infinity LC system coupled to a SCIEX QTRAP 5500 mass spectrometer. Analytes were separated on a CORTECS C18 column (2.1 x 50 mm; 1.6 µm particle size; Waters, 186007093), CORTECS C18 VanGuard Pre-column (2.1 x 5 mm, 1.6 µm particle size; Waters, 186007123) and an inline filter (0.3μM, Agilent, 5023-0271). The mobile phase gradient conditions used for chromatographic separation are detailed in Table 9. The first 0.7min and after 3.2min, the flow is directed to the waste. The mobile phase is made up of the same buffers as in targeted sugar LC-MS measurements

**Table 9:** SCFA gradient.

| **Total Time (min** | **Flow Rate (uL/min)** | **Buffer A (%)** | **Buffer B (%)** |
|---|---|---|---|
| 1.9 | 1000 | 82 | 18 |
| 2.8 | 1000 | 60 | 40 |
| 3 | 1000 | 0 | 100 |
| 4 | 1000 | 0 | 100 |
| 4.1 | 1000 | 95 | 5 |
| 5 | 1000 | 95 | 5 |

The electrospray ionization (ESI) source was set to 625 °C with a curtain gas level of 30 (arbitrary units), collision gas set to medium, an ion spray voltage of 5 500 V, and both Ion Source Gas 1 and 2 set to 90. Derivatized samples were quantified using multiple reaction monitoring (MRM).

### *In vivo* experiment

#### Animals

All mouse experiments were performed using C57BL/6 mice pre-colonized with the defined OMM12 consortium, bred and maintained under germ-free conditions at the Gnotobiotic Facility of ETH Zurich (Zurich, Switzerland). Three days prior to the start of experimental interventions, mice were transferred to positive-pressure isolator cages at the Institute of Microbiology. Animals were maintained under controlled environmental conditions with a 12-hour light/dark cycle and constant room temperature, with *ad libitum* access to sterile water and food. Animal health was monitored daily by body weight recording and clinical scoring according to established guidelines. All procedures were approved by the Cantonal Veterinary Office Zurich under license ZH092_2025.

#### Diet

Prior to dietary switches, mice were maintained on a standard autoclaved chow diet (3807 PX, Kliba Nafag; autoclaved at 134 °C). Irradiated purified diets were manufactured by Envigo Teklad (complete formulations detailed in Figure S4A). The fiber-free diet was TD.130343, formulated as previously described^10^. The XOS-supplemented diet was custom-formulated based on the TD.130343 fiber-free matrix by replacing 10% (w/w) of the glucose content with the same xylooligosaccharide substrate used in *in vitro* experiments (Biosynth, YC05381).

#### In vivo metabolomics sample preparation

Collection tubes were pre-weighed prior to sample collection and re-weighed after collecting fecal or cecal contents to determine wet sample mass. To normalize for moisture variability, a small aliquot of each sample was transferred to a pre-weighed tube and dried overnight at 60 °C; the calculated wet-to-dry mass ratio was used to normalize metabolite concentrations across samples.

For metabolite extraction, 5 mL of ice-cold MS-grade water per gram of wet tissue was added to each sample, followed by homogenization at 25 m/s for 2 min using a tissue lyser. Cellular debris and insoluble material were pelleted by centrifugation at 20,000 × g for 10 min in a pre-cooled centrifuge at 4 °C. The supernatant was transferred to a clean, pre-chilled tube, and three volumes of ice-cold MS-grade methanol were added to precipitate proteins and lipids. Samples and analytical standards were incubated at −20 °C for 1 h, followed by centrifugation at 20,000 × g for 30 min at 4 °C. The resulting supernatant was transferred to a clean pre-chilled tube, diluted, and derivatized directly for downstream measurement of short-chain fatty acids (SCFAs), polysaccharides, and monosaccharides.

### Statistical testing

Statistical analyses were performed in R using the rstatix and ggpubr packages. Data normality and homogeneity of variance were evaluated using Shapiro-Wilk and Levene’s tests, respectively. For independent group comparisons (*k* ≥ 3), normally distributed datasets with equal variance (*p* > 0.05 for both tests) were evaluated by one-way ANOVA with Tukey’s HSD post-hoc tests, whereas normal datasets violating equal variance (*p* < 0.05) were analyzed via Welch’s ANOVA with Games-Howell post-hoc tests. Non-normal datasets (*p* < 0.05) were evaluated using the Kruskal-Wallis test followed by Dunn’s post-hoc tests. For paired or repeated-measures data, normally distributed variables were assessed using one-way repeated-measures ANOVA with paired *t*-tests, while non-normal variables were analyzed using the Friedman test with paired Wilcoxon signed-rank tests. All pairwise comparisons were adjusted for multiple testing using the Benjamini-Hochberg false discovery rate (FDR) method, with significance defined at *p*_adj_ < 0.05.

## Acknowledgements

We thank all members of our laboratory for useful discussions and Vilhelmiina Haavisto for reading the manuscript. We thank Prof. Wolf-Dietrich Hardt, Manja Barthel and Dr. Bidong Nguyen for their support with the animal experiments. We would also like to thank Michal Okoniewski for his help with the transcriptomic analysis. D.L.B was supported by the Swiss National Foundation Sinergia grant (1-008812-000). N.R. and A. H. are funded by an SNSF Ambizione grant (PZ00P3_223765) to A.H. A.S. is supported through Principles of Microbial Ecosystems (PRIME) collaboration (grant 542395).

## Author contributions

Conceptualization: DLB, US

Methodology: DLB, AS, NR, AH

Investigation: DLB, NR

Visualization: DLB

Funding acquisition: US, AH

Project administration: US

Supervision: US

Writing – original draft: DLB, US

Writing – review & editing: DLB, AH, AS, US

## Declaration of generative AI and AI-assisted technologies in the writing process

During the preparation of this work, the authors used Gemini in order to increase readability and correct for language. After using this tool, the authors reviewed and edited the content as needed and take full responsibility for the content of the published article.

## Materials availability

This study did not generate new unique reagents.

## Supplementary figures

**Supplementary Figure S1:**
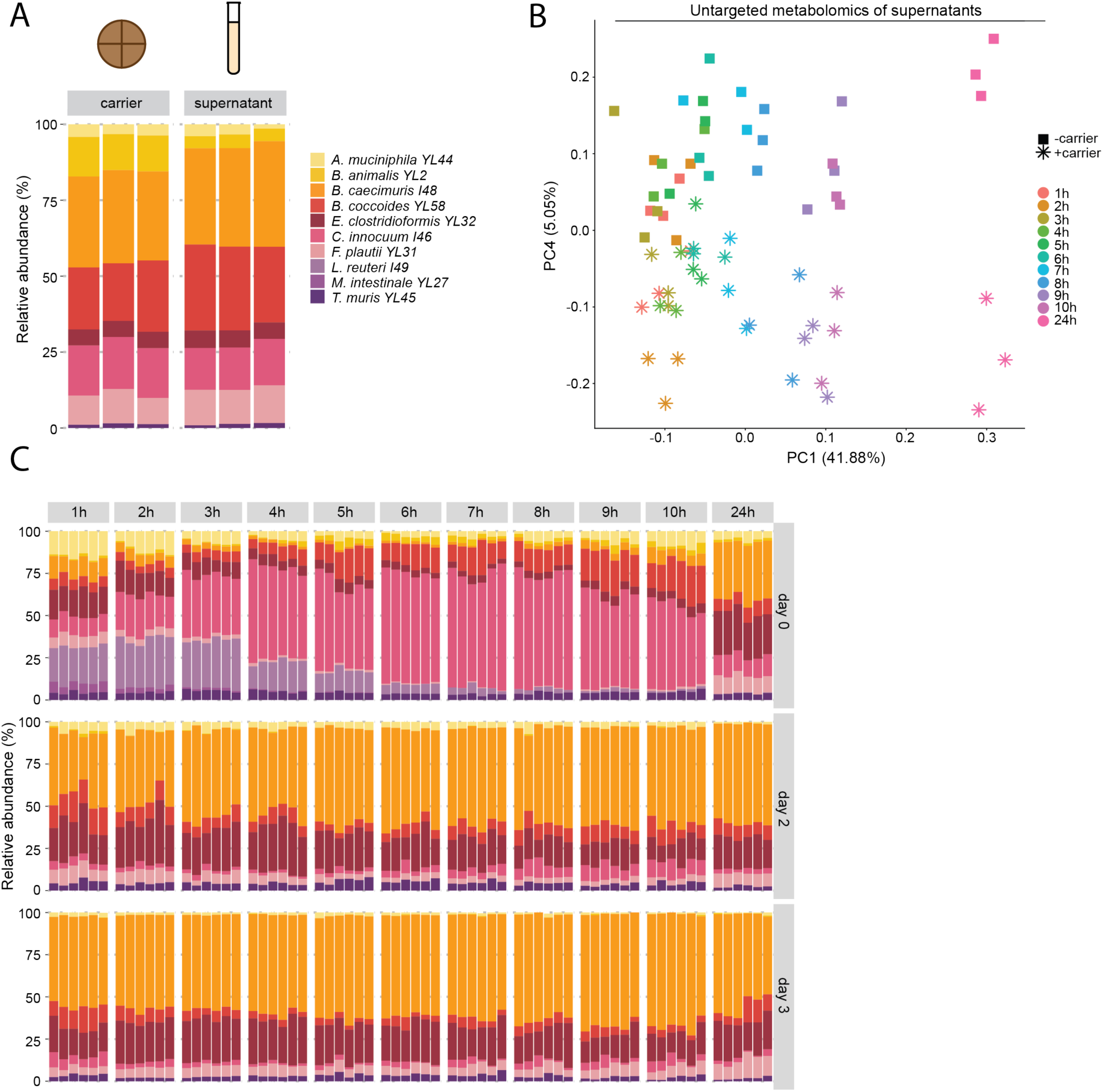
Mucin-carrier does not alter *in vitro* microbial composition or metabolome. **A)**Dynamic shifts in relative species abundance based on 16S rRNA gene profiling in APF medium on the mucin carrier vs supernatant. **B)** Principal Component Analysis (PCA) score plot of untargeted LC-MS/MS metabolomics from culture supernatants tracking metabolic trajectories across sampling time points at day 1 in the presence (+carrier, asterisks) or absence (-carrier, squares) of mucin carriers. Data are from three biological replicates per time point. **C)** Temporal dynamics of relative bacterial abundance based on 16S rRNA gene profiling during Day 0, Day 2, and Day 3 in sextuplicate cultures.

**Supplementary Figure S2:**
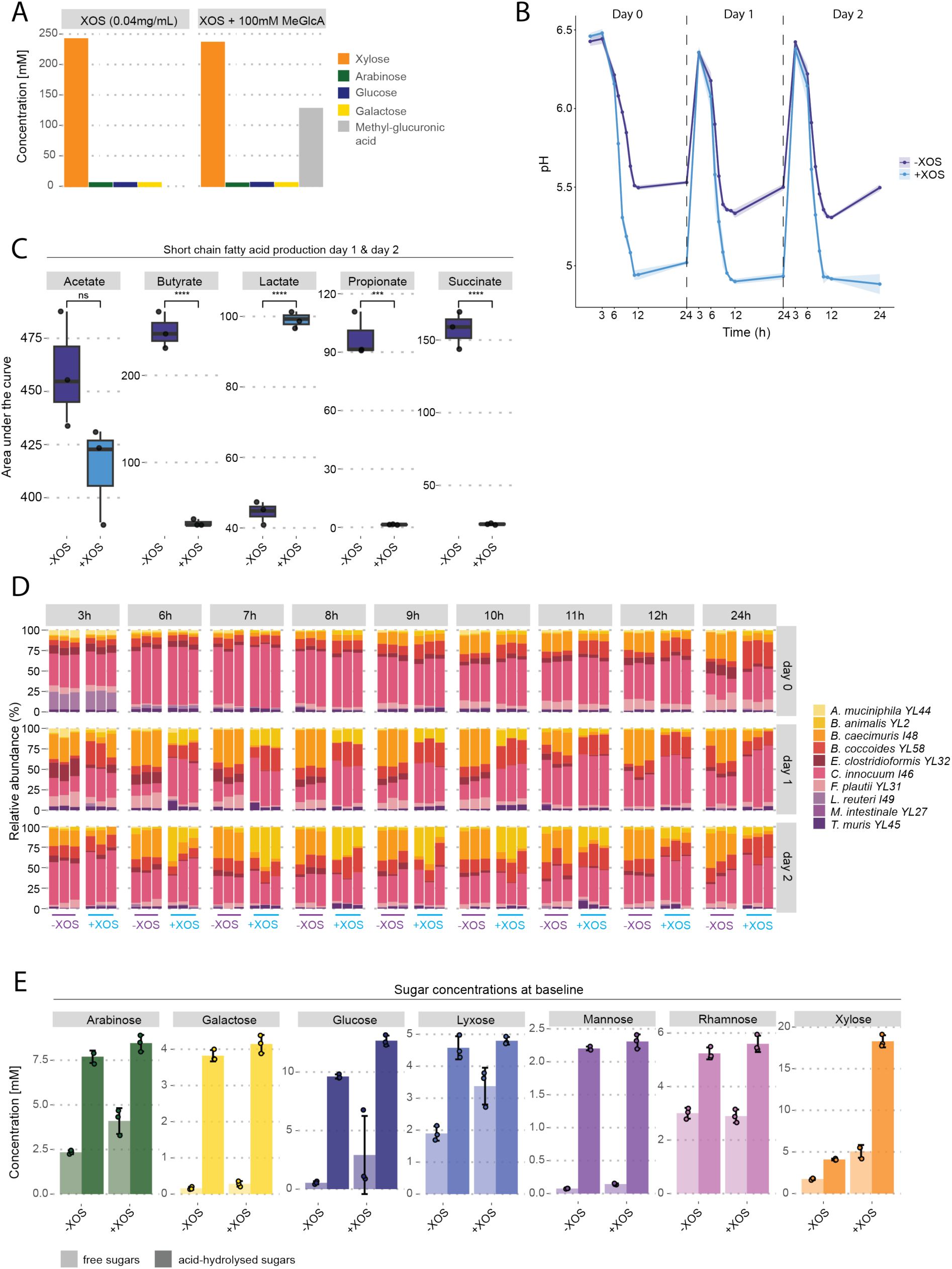
*In vitro* metabolic profiles, pH kinetics, and substrate dynamics of the OMM12 consortium. **A)** Concentration of acid-hydrolyzed monosaccharides in XOS (0.04 mg/mL) with or without 100 mM methylated glucuronic acid (MeGlcA) spike-in. **B)** Longitudinal culture supernatant pH measurements over 24-hour incubation windows across Day 0, Day 1, and Day 2 passages in control vs. XOS-supplemented conditions. Data are means ± SD from three biological replicates. **C)** Area under the curve (AUC) of short-chain fatty acid (SCFA) and lactate accumulation across consecutive passages od day 1 and day 2 in control vs. XOS-supplemented cultures. Data are means ± SD from three biological replicates. One-way ANOVA with Tukey’s post-hoc test; ***p < 0.001, ****p < 0.0001, ns = not significant. **D)** High-resolution time-course of relative bacterial abundance from 3 h to 24 h across Day 0, Day 1, and Day 2 passages in control vs. XOS-supplemented conditions. Data are from three biological replicates. **E)** Baseline concentration of free monosaccharides and acid-hydrolyzed fraction in control and XOS media. Data are means ± SD from three biological replicates.

**Supplementary Figure S3:**
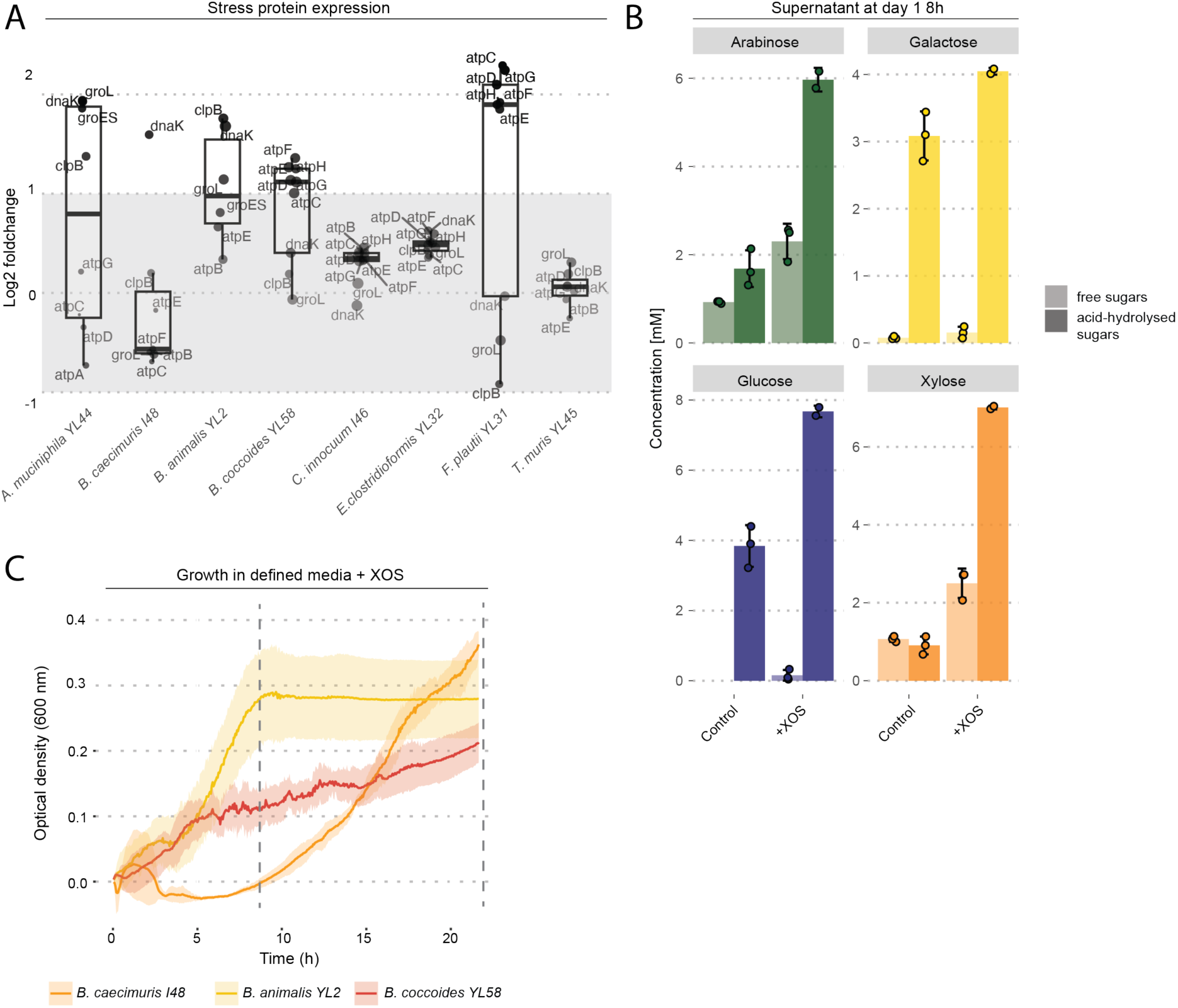
Cellular stress responses and supernatant glycan profiles at transcriptomic timepoint. **A)** Differential gene expression of molecular chaperones, stress proteins, and ATP synthase subunits across OMM12 strains in response to XOS supplementation. Boxplots show median, IQR, and range from four biological replicates. **B)** Extracellular concentrations of free and total acid-hydrolyzed monosaccharides in culture supernatants 8 hours post-inoculation on Day 1 in control vs. XOS conditions. Data are means ± SD from three biological replicates. **C)** Growth curves of primary degraders in defined media containing XOS as the sole carbon source. Dotted vertical lines represent time points chosen for sugar quantification. Data are means ± SD from three biological replicates.

**Supplementary Figure S4:**
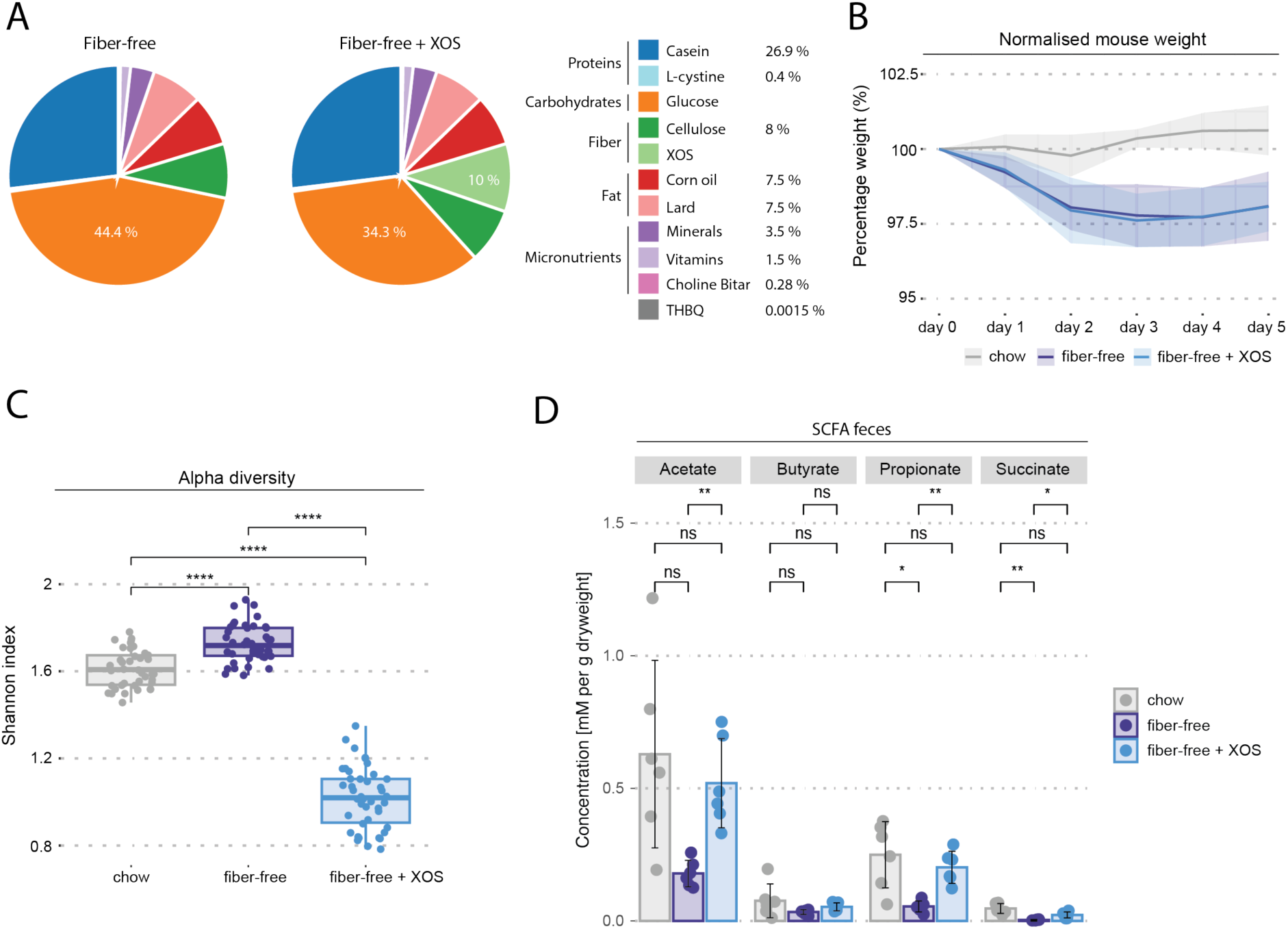
Dietary formulations, physiological parameters, and fecal short-chain fatty acids *in vivo*. **A)** Nutritional composition breakdown of experimental fiber-free and fiber-free + 10% XOS diets. **B)** Normalized body weight changes of gnotobiotic OMM12 mice over 5 days post-dietary transition from standard chow to fiber-free or fiber-free + XOS diets. Data are means ± SD of six mice per group. **C)** Fecal alpha diversity (Shannon index) across dietary cohorts across Days 1–5. Boxplots show median and IQR (N = 6 mice per group). Welch’s ANOVA with Games-Howell post-hoc test; ****p < 0.0001, ns = not significant. D) Fecal short-chain fatty acid concentrations measured on Day 6 across dietary treatment groups. Data are means ± SD (N = 6 mice per group). Welch’s ANOVA with Games-Howell post-hoc test; *p < 0.05, p < 0.01, ***p < 0.001, ns = not significant.

**Supplementary Figure S5:**
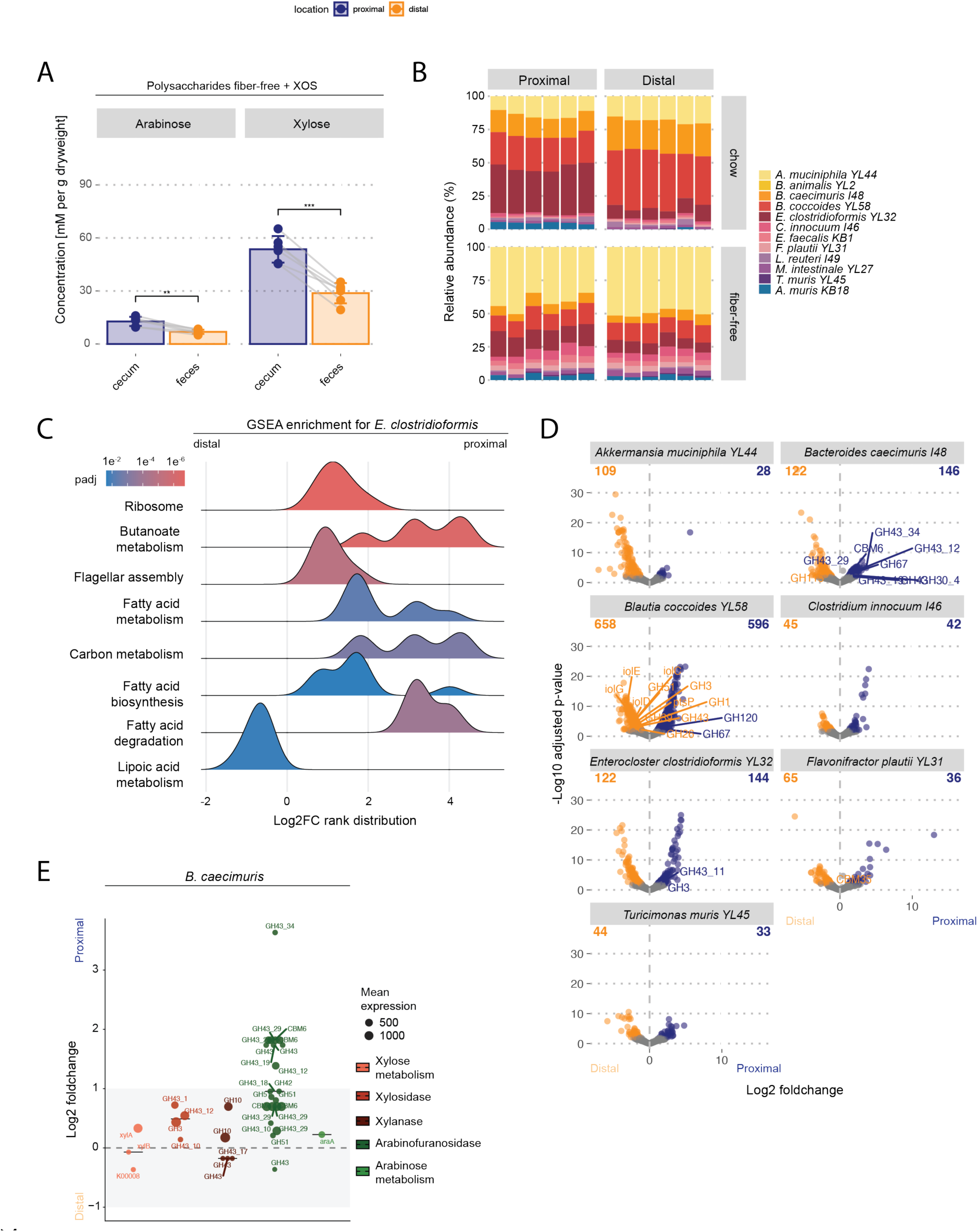
Regional carbohydrate concentration dynamics and spatial metatranscriptomics in the mouse colon. **A)** Acid-hydrolyzed polysaccharide-derived sugar concentrations in cecum and feces of XOS-supplemented mice. Data are means ± SD of six mice per group. Repeated-measures ANOVA with paired t-test; *p < 0.05, **p < 0.01. **B)** Spatial bacterial community composition based on 16S rRNA gene profiling in proximal and distal colonic segments of mice fed standard chow or fiber-free diets (N = 6 mice per group) **C)** Gene Set Enrichment Analysis (GSEA) ridge plot depicting functional pathways significantly enriched in *E. clostridioformis* between the distal and proximal colonic niches. Rank distribution is based on log2 foldchanges and the color gradient indicates adjusted p-value (padj). **D)** Strain-resolved volcano plots of spatial differential gene expression between distal and proximal colonic niches for all seven colonizing OMM12 strains in XOS-supplemented mice (N = 3 paired samples). Total upregulated DEGs indicated in upper quadrants. **E)** Spatial differential expression and baseline expression intensity (baseMean) of CAZyme genes and carbohydrate catabolic operons in *B. caecimuris* between proximal and distal colon (N = 3 paired samples). Point size scales with mean expression level.

